# Human and AI voice identities evoke shared neural signatures during speaker recognition across changes in speech content and prosody

**DOI:** 10.64898/2025.12.22.695263

**Authors:** Wenjun Chen, Marc D. Pell, Xiaoming Jiang

**Author notes:** **Corresponding author address:** Xiaoming Jiang, PhD, 1550 Wenxiang Road, Shanghai 201620, China.

## Abstract

Both biologically-produced human voices and algorithmically-generated AI speech manifest speaker identity. Critically, prosodic variations modulate the acoustic dimensions (e.g., fundamental frequency) that also shape individual speaker identity representations. So far it remains unclear whether listeners process speaker identities in human and AI voices through neurologically equivalent mechanisms, nor how prosodic cues might influence these cognitive processes. We examined event-related potentials during old/new speaker discrimination after name-based identity learning, and further analyzed correctly recognized old speakers comparing trials where prosody matched vs. mismatched between learning and testing. For old/new discrimination, multivariate pattern analysis (MVPA) revealed three significant late windows (662-1498 ms) with Pz as the primary contributor for AI voices, yet none for human voices. Univariate analyses revealed that human voices showed earlier widespread discrimination (N250: 200-280 ms), while both voice types converged on Pz as the strongest contributor based on effect size rankings for late old/new effects (400-800 ms). These old/new effects emerged across completely different speech content between learning and testing, addressing a gap in prior literature. For speaker-specific prosodic expectation effects in the 500-900 ms window, unexpected prosody elicited late positivity for human voices compared to the prosody used during learning, whereas AI voices elicited late negativity. The late positivity resembles P600 components observed for communicative style expectancy violations, while the late negativity likely reflects effortful reprocessing of prosodic violations within atypical synthetic signals, analogous to accented speech processing. Our study advances understanding of voice identity in cognitive neuroscience and offers implications for AI voices in human-computer interaction. [Word count: 250]

## 1. Introduction

Speaker identity is an essential manifestation of the self in social interaction (<u>Scott & McGettigan, 2016</u>) and our ability to rapidly identify speakers is deployed constantly: recognizing a friend’s voice within seconds of answering a phone call, distinguishing between two speakers on the radio, or immediately identifying a celebrity’s voice in an advertisement. Today, however, devices like smartphones also deliver AI-generated speech alongside human voices. Apple’s *Personal Voice* represents one such application: users record 150 sentences to clone their speaking style, allowing individuals with conditions like amyotrophic lateral sclerosis (ALS) to later “speak” by typing in FaceTime calls and assistive communication apps (<u>Apple, 2023, December 18</u>). Other systems like Huawei’s *Xiaoyi* assistant in Chinese can clone both speaker identity and prosodic style from as few as 15 sentences (<u>Chen & Jiang, 2023</u>), enabling similar usages. When the same identity exists in both human and AI voices, do listeners rely on the same perceptual mechanisms to process identity features for each voice type?

One lens to examine this question is through speaker recognition, the ability to learn and later identify individual speakers, which has revealed distinct neural mechanisms for processing familiar vs. unfamiliar voices (<u>Maguinness et al., 2018</u>; <u>Sidtis & Zäske, 2021</u>). A key motivation for examining speaker recognition across familiarity is an existing gap in understanding how identity processing generalizes when speech content or prosody varies (<u>Lavan, Knight, et al., 2019</u>; <u>Xu & Armony, 2021</u>; <u>Zäske et al., 2014</u>). We address these two gaps while using this recognition framework to compare how listeners process human vs. AI-cloned voices. Our study provides an updated understanding of the intersection of voice identity perception and how speech prosody may influence it, and most importantly, whether AI-cloned voices engage the same perceptual mechanisms as human voices.

### 1.1. Speech-content-dependent or speech-independent voice identity processing?

Voice identity arises from anatomical, acoustic, and articulatory cues and functions as a relatively stable manifestation of oneself in social communication (<u>Scott & McGettigan, 2016</u>; <u>Sidtis & Kreiman, 2012</u>). This ability is evolutionarily ancient, with newborns already showing differential neural responses to maternal vs. unfamiliar voices (<u>Adam-Darque et al., 2020</u>). Unfamiliar voices undergo a gradual learning process in which listeners repeatedly analyze their acoustic structure and compare it to an internal prototype voice (<u>Maguinness et al., 2018</u>). Through such iterative perceptual cycles, stable identity features are extracted and consolidated into a reference pattern, ultimately giving rise to the neural signatures observed for familiar vs. unfamiliar voices. A meta-analysis of functional magnetic resonance imaging (fMRI) studies revealed systematic differences across familiarity levels (<u>Sun et al., 2023</u>). Unfamiliar voices mainly activate the bilateral superior temporal gyri for basic acoustic analysis. Familiar voices additionally recruit right inferior and middle frontal gyri involved in identity processing. Higher familiarity further engages regions such as the fusiform gyrus, parahippocampal cortex, and insula, reflecting access to person-specific information.

Beyond spatial patterns revealed by fMRI, electroencephalography (EEG) studies show that voice-familiarity levels differ in their temporal dynamics. <u>Plante-Hébert et al. (2021)</u>, who manipulated familiarity through frequency of exposure rather than explicit learning phases, found that trained-familiar voices elicited a distinct N250 (300–350 ms) compared to rarely heard unfamiliar voices, whereas intimately familiar voices showed different components compared to unfamiliar voices: a P2 (200–250 ms) and a sustained LPC (450–850 ms). However, when voices are explicitly learned and later tested with manipulated speech content, different patterns emerge. <u>Zäske et al. (2014)</u> reported a parietal old/new LPC effect (300–700 ms at Pz) for correctly recognized old vs. new voices when test sentences matched study sentences, but this effect was limited to the same-sentence condition. This seemingly suggests that the LPC primarily reflects speech-content-dependent episodic retrieval rather than abstract voice representations.

Meanwhile, fMRI evidence from the same research group points to speech-independent voice representations. <u>Zäske et al. (2017)</u> used an analogous recognition paradigm but presented German sentences to non-German speakers, isolating voice processing from semantic content. Learned voices showed reduced activation compared to novel voices in right-lateralized regions including posterior superior temporal gyrus and frontal areas. Critically, unlike the LPC effect in their 2014 ERP study, these fMRI novelty reductions persisted regardless of whether test sentences matched those heard during learning, indicating that voice identity representations can generalize across speech content. These findings suggest that while fMRI reveals speech-invariant voice identity representations, current ERP markers remain sensitive primarily to speech-content-dependent episodic retrieval.

It is notable that the above studies utilized emotionally neutral prosody (<u>Plante-Hébert et al., 2021</u>; <u>Zäske et al., 2017</u>; <u>Zäske et al., 2014</u>), which minimizes cognitive load and avoids potential alterations to voice identity representations that prosodic variation might introduce (<u>Lavan, Knight, et al., 2019</u>; <u>Xu & Armony, 2021</u>). Although prior ERP studies minimized prosodic variation by using neutral prosody, they still failed to detect neural old/new differences across changes in speech content. However, there are reasons to anticipate such effects.

This expectation is grounded in behavioral work demonstrating reliable extraction of speech-invariant identity cues across utterances. In memory-intensive old/new paradigms, cross-sentence voice recognition remains robust when prosody (suprasegmental vocal features; see next section for details) is held constant: <u>Zäske et al. (2014)</u> reported accuracies of ∼62–70% across blocks using neutral prosody, and <u>Xu and Armony (2021)</u> likewise observed above-chance performance (∼67%–72%) in same-prosody/different-content trials with either fearful or neutral prosody.

Arguably, the absence of ERP differences in <u>Zäske et al. (2014)</u>, despite above-chance behavioral accuracy, reflect insufficient strength of identity encoding. This speculation is supported by evidence from explicit identity learning tasks: when listeners receive name-label training accompanied by accuracy verification, they successfully recognize speakers despite substantial within-speaker variations in glottal pulse rate (GPR, perceived as fundamental frequency, F0) and vocal tract length (VTL) that exceed those typically associated with changes in speech content (<u>Lavan, Knight, et al. (2019)</u>. Crucially, by pairing voices with names and confirming successful learning before testing, <u>Lavan, Knight, et al. (2019)</u> ensured that robust person-level representations were formed, enabling listeners to track identity-relevant acoustic variability while ignoring irrelevant within-speaker variation. By inference, implementing a similar name-labeling procedure and accuracy check should hypothetically strengthen identity encoding sufficiently to observe a parietal old/new effect at Pz as reported by <u>Zäske et al. (2014)</u> even when speech content varies, rather than merely reflecting episodic retrieval.

### 1.2. Prosody as more than acoustic variation: A speaker-specific style cue?

Beyond the linguistic content discussed above, the same sentence can be perceived quite differently depending on how it is said, that is, through speech prosody (suprasegmental features such as pitch, rhythm, and intonation patterns) (<u>Xu, 2019</u>). Prosodic variation conveys paralinguistic information about speakers’ emotional states (e.g., happy vs. sad) (<u>Pell & Skorup, 2008</u>), epistemic certainty (e.g., feeling of knowing: confident vs. doubtful) (<u>Jiang & Pell, 2015</u>; <u>Jiang & Pell, 2017</u>), and communicative intentions (e.g., sincere vs. ironic) (<u>Fish et al., 2017</u>; <u>Mauchand et al., 2020</u>).

While variation in speech content with similar prosodic patterns likely induces only small within-speaker fluctuations in structural identity cues, such as F0 and VTL (<u>Mathias & von Kriegstein, 2019</u>), changes in paralinguistic cues can be far more disruptive (<u>Anikin et al., 2021</u>; <u>Belyk et al., 2022</u>; <u>Pisanski et al., 2022</u>). Real-time MRI studies show that emotional prosody systematically shifts vocal-tract configuration, with happy speech produced using shorter effective VTL and greater articulatory opening than angry, sad, or neutral speech (<u>Kim et al., 2020</u>). Meanwhile, F0 and VTL function in an inverse relationship when listeners estimate vocal-tract dimensions (<u>Darwin et al., 2003</u>; <u>Neuhaus et al., 2024</u>). Prosodic variations across emotional, social, and pragmatic contexts are thus characterized by systematic modulations of both F0 and VTL (<u>Larrouy-Maestri et al., 2025</u>). For instance, confident prosody is produced with lower F0 and longer effective VTL than doubtful prosody in both English (<u>Jiang & Pell, 2017</u>) and Mandarin (<u>Chen & Jiang, 2023</u>). Most importantly, listeners are sensitive to such coordinated shifts in VTL and F0, influencing how they derive individual identity representations (<u>Lavan, Burston, et al., 2019</u>; <u>Lavan, Knight, et al., 2019</u>).

Given that prosodic shifts systematically modulate the same acoustic dimensions that define speaker identity (<u>Lavan, Burton, et al., 2019</u>; <u>Pinheiro, 2025</u>), a critical question arises: do listeners treat prosody merely as acoustic variability to be normalized, or do they encode prosodic patterns as part of a speaker’s identity representation? Evidence from syntactic processing suggests the latter may be plausible. <u>Kroczek and Gunter (2021)</u> showed that listeners internalize a speaker’s habitual syntactic preferences (e.g., marked OSV vs. SVO word order), and violations of these speaker-specific patterns elicit P600 responses even when sentence meaning remains unchanged. This demonstrates that listeners bind linguistic style to speaker identity.

We propose that prosodic style may function analogously. If listeners learn a talker under one prosodic style but later encounter a different style from the same speaker, this mismatch may elicit expectancy-violation responses. Such findings would carry two implications. First, prosodic style, like syntactic style, functions as a speaker-specific cue bound to identity (<u>Kroczek & Gunter, 2021</u>). Second, successful recognition despite prosodic shifts would indicate that listeners extract identity representations that generalize across acoustic variation, with P600-like prediction errors (<u>Kroczek & Gunter, 2021</u>) signaling the comparison between stored and incoming identity cues (<u>Lavan, Burston, et al., 2019</u>; <u>Lavan, Knight, et al., 2019</u>).

### 1.3. Are AI and human speaker identities represented similarly in the cognitive system?

Human voice production relies on physiological articulatory systems, whereas AI-synthesized speech is generated by deep learning models (<u>Khanjani et al., 2023</u>), with acoustic features derived from parametric structures rather than anatomical constraints. Thus, the frameworks discussed in the preceding subsections concerning how speech content and prosody influence identity recognition are grounded in human voice perception and cannot be automatically generalized to AI voices. Currently, there exists no systematic understanding of whether AI voices support identity memorization and recognition at behavioral and neural levels. Based on existing relevant evidence, we propose that AI voice identity learning and recognition mechanisms may share components with human voice perception.

One source of evidence comes from deepfake detection tasks, predominantly from English-language studies, where listeners judge whether voices originate from AI or humans. Large-scale studies report variable human performance. <u>Warren et al. (2024)</u> found overall accuracies of 63.9–85.8% across three benchmark datasets with over 1,200 participants, while <u>Müller et al. (2022)</u> reported that human listeners’ accuracy plateaued at about 80% in a gamified ASVspoof2019 detection task (410 participants, 13,229 trials). More recently, <u>Barrington et al. (2025)</u> employed state-of-the-art *ElevenLabs* voice cloning and showed that listeners correctly identified AI-generated voices as synthetic on only about 60% of trials, indicating that high-quality voice clones frequently evade explicit human detection. These behavioral findings suggest that although humans can detect degraded synthetic speech, high-quality AI clones often pass as human.

Importantly, neuroimaging work demonstrates dissociable neural pathways for natural and synthetic identities. <u>Roswandowitz et al. (2024)</u> reported that despite above-chance deepfake identity matching (∼69%), natural and deepfake voices engaged distinct neural pathways: natural identities activated the nucleus accumbens (NAcc), a region linked to social reward and bonding, whereas deepfake voices showed weaker NAcc responses but stronger auditory-cortical activity, suggesting synthetic voices lack socially rewarding properties and require greater acoustic-phonetic processing effort. Similarly, <u>Bratan et al. (2025)</u> found that AI-synthesized emotional voices, though judged acoustically similar to natural voices, elicited substantially weaker activation in mirror-neuron and emotion-related regions, instead primarily recruiting memory-related networks. These neural dissociations imply that despite partial behavioral deception, synthetic voices are not processed as fully natural speaker identities by the human brain.

A second source of evidence comes from research directly exploring identity representation. In identity matching tasks where listeners judge whether two consecutive audio clips are the same speaker, <u>Barrington et al. (2025)</u> found that participants judged AI-cloned voices and their human sources as the same identity in approximately 80% of trials, indicating that listeners often treat cloned and original voices as belonging to a single speaker. However, <u>Chen et al. (2025)</u> used AI-generated voices whose human-likeness had been independently validated as substantially lower than that of natural voices. Under these conditions, listeners judged AI clones and their human sources as the same identity only slightly above chance level, suggesting that awareness of voice source matters for identity perception. <u>Chen et al. (2025)</u> also showed that within AI or human voices separately, identity recognition accuracy reached a 99% ceiling when prosody was consistent and remained above 90% when prosody differed. This indicates that both human and AI voices carry internally coherent identity signatures to which listeners are highly sensitive. We therefore propose that when AI and human voices are examined separately, without requiring cross-category identity recognition, both groups should exhibit broadly similar behavioral patterns of identity discrimination, albeit with lower accuracy under memory-demanding recognition tasks than in short-term matching paradigms.

Beyond cross-category comparisons between AI and human voices, a third line of evidence examines whether AI voices can engage person-specific representation systems when identity familiarity is manipulated within synthetic speech alone. Using functional near-infrared spectroscopy (fNIRS), <u>Zhang et al. (2025)</u> showed that AI-generated maternal voices, cloned from participants’ own mothers, elicited significantly stronger activation in listeners’ prefrontal and temporal cortices than AI-generated unfamiliar female voices, consistent with engagement of familiarity-, emotion-, and memory-related processes. Although this study did not compare AI against natural speech or test identity recognition per se, it demonstrates that person-specific familiarity signals can be reinstated even in fully synthetic speech, supporting the idea that AI voices may access overlapping person-representation systems.

Taken together, existing evidence shows that AI voices can indeed deceive listeners into perceiving them as human and can engage familiarity-based identity processing to some extent, but their neural signatures could still remain distinct from those of natural voices in certain aspects.

### 1.4. The present study

As demonstrated above, existing studies have not simultaneously examined speaker recognition in the face of both speech content and prosodic variations, nor have they directly compared the neural mechanisms of identity processing between AI and human voices. The present study addresses these gaps by comparing identity learning and recognition in human vs. AI voices across manipulations of speech content and prosody. We employ AI clones with prosodic styles cloned separately to minimize acoustic confounds (<u>Roswandowitz et al., 2024</u>). Human and AI voices are examined in separate blocks. Participants learn speaker identities through name-labeling, then perform old/new recognition with different sentences while prosodic consistency is manipulated.

To examine these questions, we combine multivariate pattern analysis (MVPA) to identify when neural patterns discriminate conditions with univariate ERP analyses to characterize established voice recognition components. We propose three sets of predictions:

- ***Speech-content-independent voice recognition***. Parietal old/new effects should emerge across changes in speech content when prosody remains consistent (<u>Xu & Armony, 2021</u>; <u>Zäske et al., 2014</u>). Additionally, we anticipate replicating the speech-independent beta band oscillations (16-17 Hz, 290-370 ms) at central and right temporal sites reported by <u>Zäske et al. (2014)</u>, which may reflect content-invariant voice identity processing distinct from episodic memory retrieval.
- ***Speaker-specific prosody expectation effect***. Prosodic violations should produce late ERP components reflecting speaker-specific expectancy violations analogous to syntactic coupling effects (<u>Kroczek & Gunter, 2021</u>).
- ***AI-human comparability***. We anticipate that both voice types should show parietal old/new effects given comparable within-category identity discrimination (<u>Chen et al., 2024</u>). Both should also exhibit speaker-specific prosody expectation effects. However, given that natural voices engage socio-emotional and embodied-resonance networks more strongly than synthetic voices (<u>Bratan et al., 2025</u>; <u>Roswandowitz et al., 2024</u>), we predict that prosodic violation responses in human voices should resemble P600-like expectancy violations (<u>Kroczek & Gunter, 2021</u>). In contrast, AI voices may show nuanced differences in spatial distributions, component polarities, or latencies, though we cannot specify with certainty.

## 2. Methods

### 2.1. Participants

We recruited 43 native Mandarin speakers from universities in Shanghai. Three participants were excluded due to technical issues in the initial experimental setup. The remaining 40 participants consisted of 20 females (age: 21.7 ± 1.7 years; years of education: 17.6 ± 1.9) and 20 males (age: 23.2 ± 1.7 years; years of education: 18.8 ± 1.7). None reported speech or hearing impairments or any psychiatric or neurological disorders. All provided written informed consent, and the study was approved by the Ethics Committee of the Institute of Language Sciences at Shanghai International Studies University. Participants received 50 RMB per hour as compensation. Our sample size of 40 participants is consistent with established EEG studies of speaker recognition, such as <u>Zäske et al. (2014)</u> who tested 24 participants.

### 2.2. Stimuli

#### 2.2.1. Stimuli creation

Our audio stimuli were selected from a larger validated corpus of 11,808 audio recordings. The large corpus was created through a multi-stage procedure: First, 24 native Mandarin speakers (12 females, 12 males) recorded 15 sentences in three prosodic styles (confident, doubtful, neutral), which were used to train speaker-specific AI voice models using Huawei’s *Celia* system; part of the acoustic validation for 10 of these speakers is reported in <u>Chen and Jiang (2023)</u>.

Detailed procedures for Steps 2-4 are reported in (<u>Chen et al., 2025</u>). Second, these AI models generated 123 novel sentences in confident and doubtful prosodies (5,904 AI recordings: 123 sentences × 2 prosodies × 24 speakers). Critically, each speaker’s confident AI clone was trained exclusively on their confident recordings, and their doubtful AI clone exclusively on their doubtful recordings, with no cross-training between prosodic styles. Third, approximately one month later, the same 24 speakers returned to produce human versions of these 123 sentences in both prosodic styles, resulting in 5,904 human recordings matched to the AI corpus. Fourth, 48 independent listeners rated 11,808 clips (human and AI, confident and doubtful) on perceived humanlikeness and vocal confidence using 7-point scales.

For the present EEG study, 960 stimuli were selected based on these validation ratings. Sentences were brief Mandarin Chinese statements expressing factual information (e.g., [My mouse is broken]), evaluations (e.g., [She has a good sense of humor]), or intentions (e.g., [They don’t want to go to class anymore]).

The 24 speakers were divided into four groups of six by selecting every other speaker from height-ranked sequences within each biological sex (e.g., speakers ranked 1st, 3rd, 5th, 7th, 9th, 11th formed one group), creating perceptual similarity gradients while controlling for height-related acoustic cues (F0). Within each group, stimuli were selected based on validation ratings to ensure that confident prosody received higher confidence ratings than doubtful prosody, and human voices received higher humanlikeness ratings than AI voices (mean differences, not statistically tested).

The audio files were normalized to –30 dBFS and standardized to stereo format (44,100 Hz sampling rate) using *pydub* library (version 0.25.1) in Python 3.11.4 (Robert, 2021) to ensure that all stimuli were presented at the same volume level through headphones during EEG recording.

#### 2.2.2. Stimuli validation

We conducted acoustic and perceptual validation on all 960 voice recordings (480 human-AI speaker pairs). F0 was extracted using *Praat* 6.2.09 (<u>Boersma & Weenink, 2021</u>). Mean F0 was computed for each utterance after using the *extractvowels* plugin from the *Praat Vocal Toolkit* to extract each utterance’s vowels (<u>Chen et al., 2025</u>; <u>Corretgé, 2024</u>). Independent raters evaluated all stimuli on two 7-point scales: confidence level (1 = not confident at all, 7 = very confident) and humanlikeness (1 = very machine-like, 7 = very human-like). Statistical analyses used Linear Mixed-Effects Regression (LMER) implemented in R (version 4.3.3) with the *lmerTest* package (<u>Kuznetsova et al., 2017</u>). For F0, the model formula was: `f0_mean_hz ∼ Source * Confidence_level * Biological_Sex + (1|Speaker) + (1|Item)`. For perceptual ratings, model formulas were: `perceived_humanlikeness ∼ Source * Confidence_level + (1|Speaker) + (1|Item)` and `perceived_confidence ∼ Source * Confidence_level + (1|Speaker) + (1|Item)`. Simple effects were examined using the *emmeans* package (<u>Lenth et al., 2021</u>), and effect sizes were quantified using Cohen’s *d*.

Our validation analyses confirmed successful experimental manipulations. For F0 (**Figure 1A**), human voices differed from AI voices in the doubtful condition (*p <* 0.001, *d* = 0.09) but not in the confident condition (*p =* 0.146, *d* = –0.03). Prosody effects were significant for both human (*p <* 0.001, *d* = 0.26) and AI voices (*p <* 0.001, *d* = 0.14). For confidence ratings (**Figure 1B**; left panel), human voices were rated higher than AI voices in both prosody conditions (confident: *p <* 0.001, *d* = 0.84; doubtful: *p <* 0.001, *d* = 1.07). Prosody effects on confidence ratings were significant for both human (*p <* 0.001, *d* = 5.95) and AI voices (*p <* 0.001, *d* = 2.54). For humanlikeness ratings (**Figure 1B**; right panel), human voices were rated higher than AI voices in both confident (*p <* 0.001, *d* = 4.42) and doubtful (*p <* 0.001, *d* = 4.56) conditions. Prosody effects on humanlikeness were non-significant for human voices (*p =* 0.056, *d* = 0.24) but significant for AI voices (*p <* 0.001, *d* = 0.31).

**Figure 1.**
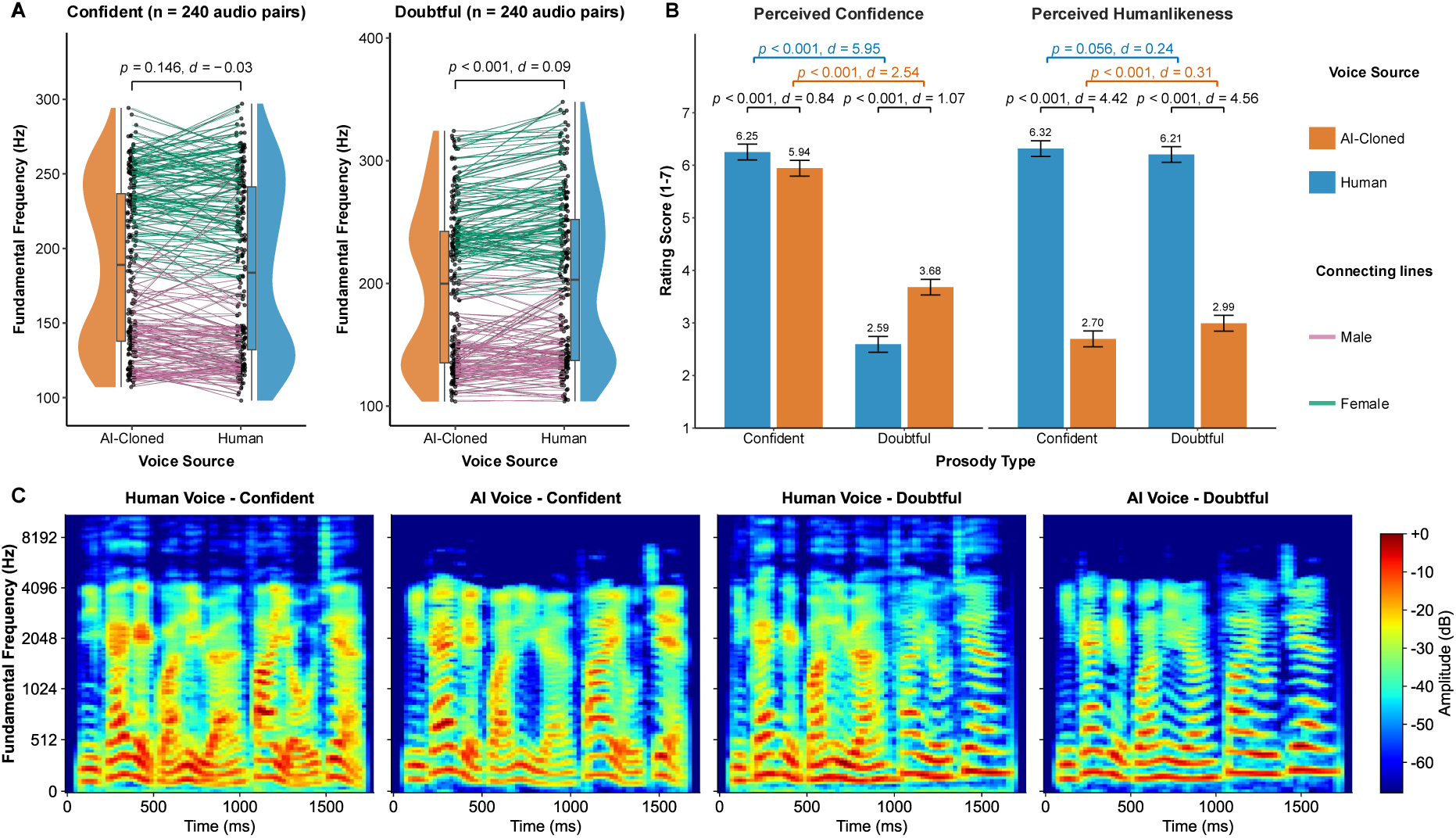
Acoustic and perceptual characteristics of 960 voice stimuli used in the EEG experiment. *Note*. **(A)** F0 distributions for human and AI-cloned voices across prosodic conditions. Connecting lines link matched human-AI speaker pairs, colored by male/female. **(B)** Perceptual ratings showing perceived confidence and humanlikeness across voice sources and prosody types. **(C)** Example of one male speaker producing the Mandarin sentence [You should send an email to the administrator]. AI voices: generated by independently trained models (confident clone trained on confident speech only; doubtful clone trained on doubtful speech only). Human voices: recorded approximately one month later, with the speaker instructed to naturally produce the sentence after hearing the AI-generated version. Color intensity represents amplitude (dB).

To visualize acoustic patterns, we generated mel spectrograms for one representative human-AI matched speaker pair across both prosody conditions using the *librosa* package (version 0.11.0; (<u>McFee et al., 2025</u>)) in Python 3.14. Spectrograms were computed with the following parameters: sampling rate = 22,050 Hz, FFT window size = 2,048 samples, hop length = 512 samples, and 128 mel frequency bands. Power spectrograms were converted to decibel scale relative to peak amplitude. To enable direct visual comparison of acoustic patterns across utterances,, we standardized leading and trailing silence to 80 ms for all displayed utterances using the *pydub* library (version 0.25.1) (<u>Robert, 2021</u>). **Figure 1C** shows an example mel spectrogram comparison for one male speaker-clone pair.

### 2.3. Experiment details

#### Experimental design

Participants completed eight blocks with fixed voice source order (human blocks 1-4, AI blocks 5-8). This fixed ordering was implemented because we could not assume uniform participant responses to alternating between biologically-produced and algorithmically-generated voices across blocks, which could introduce variable task-switching costs and compromise data consistency during identity learning. This design choice is justified by our behavioral results (see Checking phase accuracy and 4.3 and 4.4 in Discussion). Meanwhile, prosody order was counterbalanced across participants: half (n=20; 10 males, 10 females) experienced confident-then-doubtful prosody (Blocks 1-2, 5-6 confident; Blocks 3-4, 7-8 doubtful), while the other half (n=20; 10 males, 10 females) experienced doubtful-then-confident prosody. Each block lasted approximately 10 minutes, with the entire experiment taking 90-100 minutes including breaks.

#### Procedure

The experiment consisted of eight blocks (human and AI voices never mixed within blocks), each comprising three phases: Training, Checking, and Testing (**Figure 2A**). During Training (24 trials), participants learned to associate three unfamiliar speakers’ voices with Chinese surnames (e.g., [Junior ZHAO]). In the Checking phase (12 trials), participants identified speakers using a three-alternative forced-choice task to verify successful identity learning. During Testing (72 trials), participants categorized speakers as OLD (trained) or NEW (untrained). Training and Testing used completely different linguistic materials (sentences). The key manipulation was prosody consistency: participants learned speaker identities through voices in either confident or doubtful prosody during Training but encountered these speakers in both the same and different prosodic conditions during Testing (**Figure 2B-C**). Participants proceeded at their own pace during Training to encode speaker-name associations, but were instructed to respond quickly and accurately during Checking and Testing phases. An accuracy threshold of 10/12 correct responses (≥83%) was required in the Checking phase; accuracy feedback was displayed after each block, and blocks failing to meet this threshold were repeated after a brief break.

**Figure 2.**
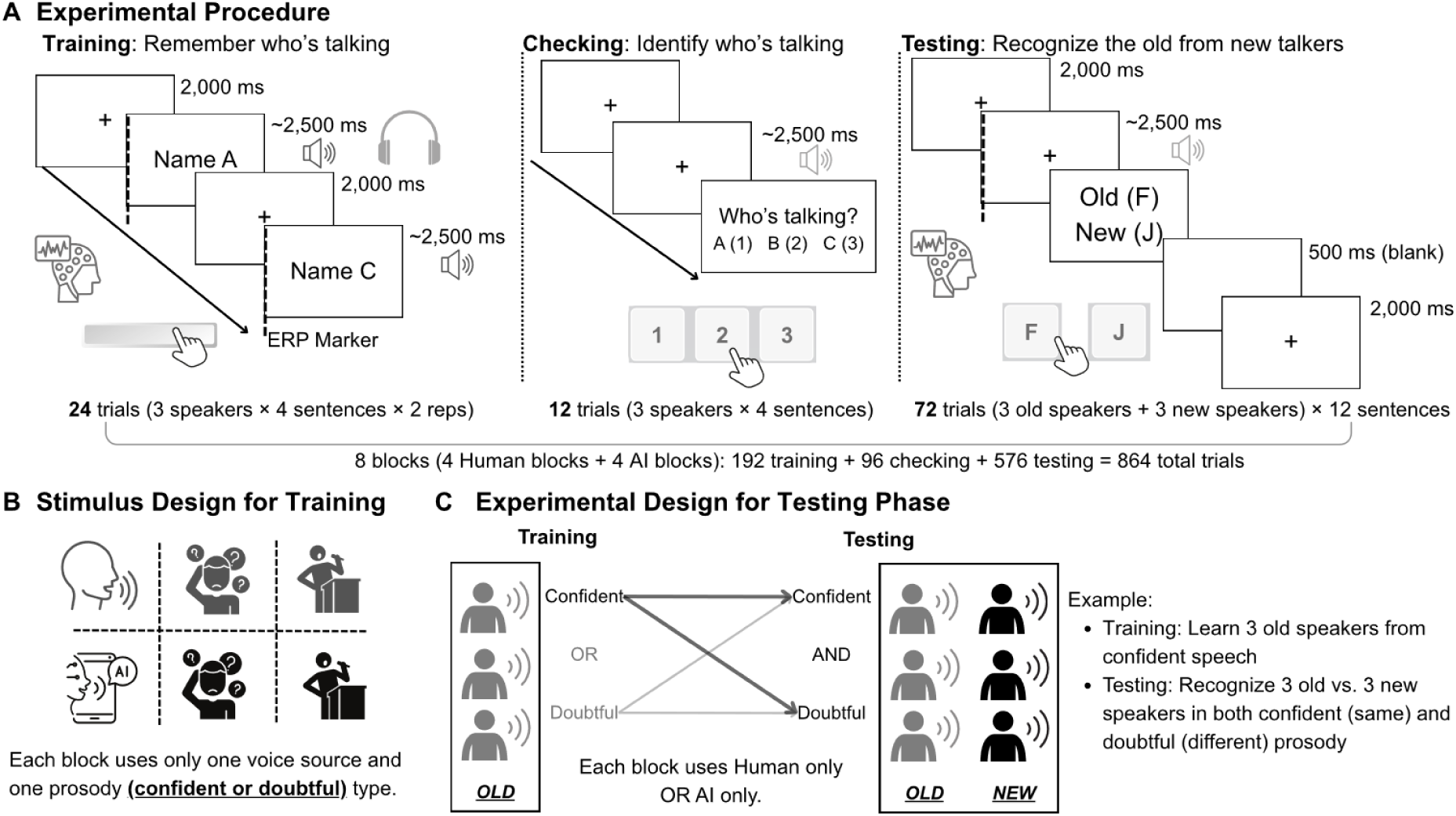
Experimental design and procedure for speaker identity learning and recognition. *Note.* **(A)** Each of 8 blocks included three phases: Training, where participants learned three speakers’ names from their voices (24 trials); Checking, where participants identified speakers using the same utterances (12 trials, three-alternative forced-choice); and Testing, where participants categorized speakers as OLD (trained) or NEW (untrained) (72 trials). **(B)** Training stimulus design showing that each block used one voice source (Human or AI) and one prosody (confident or doubtful). **(C)** Testing design where participants distinguished OLD from NEW speakers, with prosody consistency (same vs. different from Training) as the critical manipulation.

#### EEG recording

EEG data were acquired in a sound-treated, dimly lit electromagnetically shielded laboratory. Continuous EEG was recorded using a 64-channel elastic cap with passive Ag/AgCl electrodes (ActiCap system, Brain Products, Germany) according to the extended 10-20 system. FCz served as the online reference electrode.

Electrode impedances were maintained below 5 kΩ using conductive gel. Signals were digitized at 500 Hz with a bandpass filter of 0.01-100 Hz. Two electrooculography electrodes were placed near the left outer canthus and below the right eye to monitor horizontal and vertical eye movements and blinks. Auditory stimuli were presented binaurally through audiology insert earphones. Participants sat in a comfortable chair approximately 80 cm from a computer monitor in the shielded room. Participants were instructed to minimize head and body movements and to blink only between trials when possible.

### 2.4. Data analysis

#### Behavioral data analysis

Behavioral performance was analyzed using LMERs implemented in R (version 4.3.3) with the *lmerTest* package (<u>Kuznetsova et al., 2017</u>). The Checking phase aimed to ensure that participants successfully established associations between speaker identities and their voices. Two separate models were fitted for accuracy and reaction time (RT), respectively: `accuracy ∼ Source * Prosody + (1|Participant)` and `rt ∼ Source * Prosody + (1|Participant)`, where Source (human vs. AI) and Prosody (confident vs. doubtful) were fixed effects. Post-hoc pairwise comparisons were conducted using the *emmeans* package (<u>Lenth et al., 2021</u>) to compare prosody conditions within each voice source. Effect sizes were calculated using Cohen’s *d* with pooled standard deviations.

For the Testing phase (Old/New effect), two sets of analyses were conducted. The first analysis examined old/new speaker recognition effects: `accuracy ∼ Source * Old/New + (1|Participant)` and `rt ∼ Source * Old/New + (1|Participant)`, where Old/New compared old vs. new speakers. For RT analysis here, data were filtered to include only correct trials.

For the prosody expectation effect, only old speaker trials were retained. The analysis used the formula: `accuracy ∼ Source * Prosody + (1|Participant)` and `rt ∼ Source * Prosody + (1|Participant)`, where Prosody compared same vs. different prosody relative to the Training phase. For RT analysis, data were further filtered to include only trials with correct same/different judgments. Post-hoc comparisons and effect size calculations followed the same procedures as the Checking phase.

RT analyses retained correct trials only. For Old/New recognition, retention rates were 62.2% for Human Old voices (3,584 of 5,760 trials), 74.4% for Human New voices (4,283 of 5,760 trials), 60.5% for AI Old voices (3,484 of 5,760 trials), and 73.5% for AI New voices (4,232 of 5,760 trials). For the prosody expectation effect analyses on old trials, retention rates in Same prosody conditions were high for both Human (74.7%, 2,152 of 2,880 trials) and AI voices (74.5%, 2,145 of 2,880 trials). However, retention rates dropped in Different prosody conditions for both Human (49.7%, 1,432 of 2,880 trials) and AI voices (46.5%, 1,339 of 2,880 trials).

#### EEG data preprocessing

The preprocessing was conducted using EEGLAB (version 2025.0) (<u>Delorme & Makeig, 2004</u>) in MATLAB (version R2024b). Data were re-referenced offline to the average of bilateral mastoid electrodes. To account for leading silence differences between manually edited human recordings and AI-generated audio, silence duration was quantified for each file using the *pydub* library (version 0.25.1) (<u>Robert, 2021</u>) in Python 3.11.7, identifying silence below –50 dB lasting greater than 5 ms. EEG event markers were then adjusted by these durations to ensure precise time-locking to acoustic onset. Raw continuous EEG data were band-pass filtered. Bad channels and trials with obvious artifacts were identified through visual inspection; bad channels were interpolated using spherical spline interpolation, and contaminated trials were removed. For ERP analysis, data were filtered at 0.1–40 Hz, epoched from –500 to 1500 ms relative to the adjusted stimulus onset, and baseline-corrected using the –500 to 0 ms pre-stimulus interval.

For time-frequency analysis, data were filtered at 0.1–100 Hz, epoched from –900 to 1500 ms, and baseline-corrected using the –500 to –250 ms interval. For Independent Component Analysis (ICA), data were temporarily high-pass filtered at 1 Hz to improve decomposition quality. ICA decomposition using the extended infomax algorithm was performed on the 1 Hz filtered data, and the resulting unmixing weights were transferred back to the 0.1 Hz filtered data. Components reflecting ocular, muscular, and cardiac artifacts were identified through visual inspection of component topographies and time courses, then removed by back-projection. After artifact removal, data were low-pass filtered at 40 Hz for ERP analysis and 100 Hz for time-frequency analysis.

#### MVPA analysis

To identify when neural activity patterns discriminated between experimental conditions, we applied MVPA to the time-domain EEG data using linear discriminant analysis (LDA) implemented in the MVPA-Light toolbox for MATLAB (<u>Treder, 2020</u>). ERP epochs (−500 to 1500 ms relative to stimulus onset) were analyzed at each time point, with a classifier trained using the spatial pattern of voltage across all electrodes. Two decoding analyses were performed: (1) distinguishing old vs. new speakers (correct trials only), and (2) distinguishing same vs. different prosody within correctly recognized old speakers. Classification performance was evaluated using 5-fold cross-validation, with decoding accuracy quantified using the area under the receiver operating characteristic curve (AUC) and classification accuracy. This procedure was applied separately for human and AI voices, yielding time-resolved decoding trajectories for each participant. Group-level statistical inference was performed using cluster-based permutation testing (1,000 permutations) to control for multiple comparisons across time. Significant clusters were identified using a cluster-forming threshold of *z* = 1.96 (corresponding to *p* < 0.05), with a family-wise error rate of *α* = 0.05, as contiguous temporal windows where decoding performance exceeded chance level (AUC = 0.5).

Additionally, time-frequency MVPA was conducted to examine oscillatory power patterns. Given the absence of statistically significant effects, these results are reported in **Supplementary Analysis 1**.

#### Old-new effect (MVPA-based clusters)

Based on the temporal MVPA results, three significant time windows were identified for AI voices in the old vs. new speaker recognition task. Direct MVPA on human voices did not identify significant clusters, likely because human voices contain inherently greater acoustic variability than AI voices (**Figure 1C**), which may have obscured identity-based discrimination signals. We therefore applied these AI-derived time windows to human voice data for comparative analysis. For each cluster, we identified the top 10 electrodes with the highest contribution to decoding accuracy using Haufe-transformed LDA weights (<u>Haufe et al., 2014</u>). Trial-level ERP amplitudes averaged across these 10 electrodes within each time window were analyzed using LMERs (<u>Kuznetsova et al., 2017</u>): `Voltage ∼ OldNew + (1|Participant) + (1|Item).` Type III ANOVA with Satterthwaite’s approximation tested the main effect of speaker identity (old vs. new), with effect sizes quantified using partial omega-squared (ω²).

#### Old-new effect (Literature-based)

To examine condition-related amplitude differences across scalp regions, we conducted LMER analyses on trial-level ERP data extracted from nine electrodes (F3, Fz, F4, C3, Cz, C4, P3, Pz, P4) selected based on prior voice recognition research (<u>Zäske et al., 2014</u>). For each electrode, we averaged voltage values within three predefined time windows: N250 (200-280 ms), P3 (300-380 ms), and LPC (400-800 ms) (<u>Plante-Hébert et al., 2021</u>). The same LMER modeling approach was used with statistical inference via Type III ANOVA (Satterthwaite approximation) and effect size quantification via ω². Pairwise contrasts between old and new conditions were conducted using estimated marginal means with the *emmeans* package (<u>Lenth et al., 2021</u>). The alpha level was set at .05 for all tests.

#### Speaker-specific cue-binding effect (Literature-based)

To investigate whether prosodic consistency between learning and test phases influenced recognition performance, we conducted LMER analyses examining the same vs. different prosody contrast within correctly recognized old speakers. Based on previous research demonstrating late ERP effects for speaker-specific expectancy violations (<u>Kroczek & Gunter, 2021</u>), we analyzed a late time window (500-900 ms). Trial-level ERP amplitudes averaged across these electrodes were analyzed using the model formula `Voltage ∼ SameDiff + (1|Participant) + (1|Item)`, testing the main effect of prosodic consistency. All other statistical procedures remained identical to the old vs. new analysis.

## 3. Results

### 3.1. Behavioral results

In the Training/Checking phase, participants heard each audio clip twice and then identified the correct name from three options after hearing a single audio presentation. For human voices, we found neither accuracy nor reaction time differed between prosody conditions. In contrast, for AI voices, participants achieved higher accuracy (*z* = –2.94, *p* = .003, *d* = 0.61) and faster responses (*z* = 4.40, *p* < .001, *d* = 0.55) when learning speakers with doubtful prosody (94.7%, 1167 ms) compared to confident prosody (91.6%, 1514 ms). See **Figure 3A-B**.

**Figure 3.**
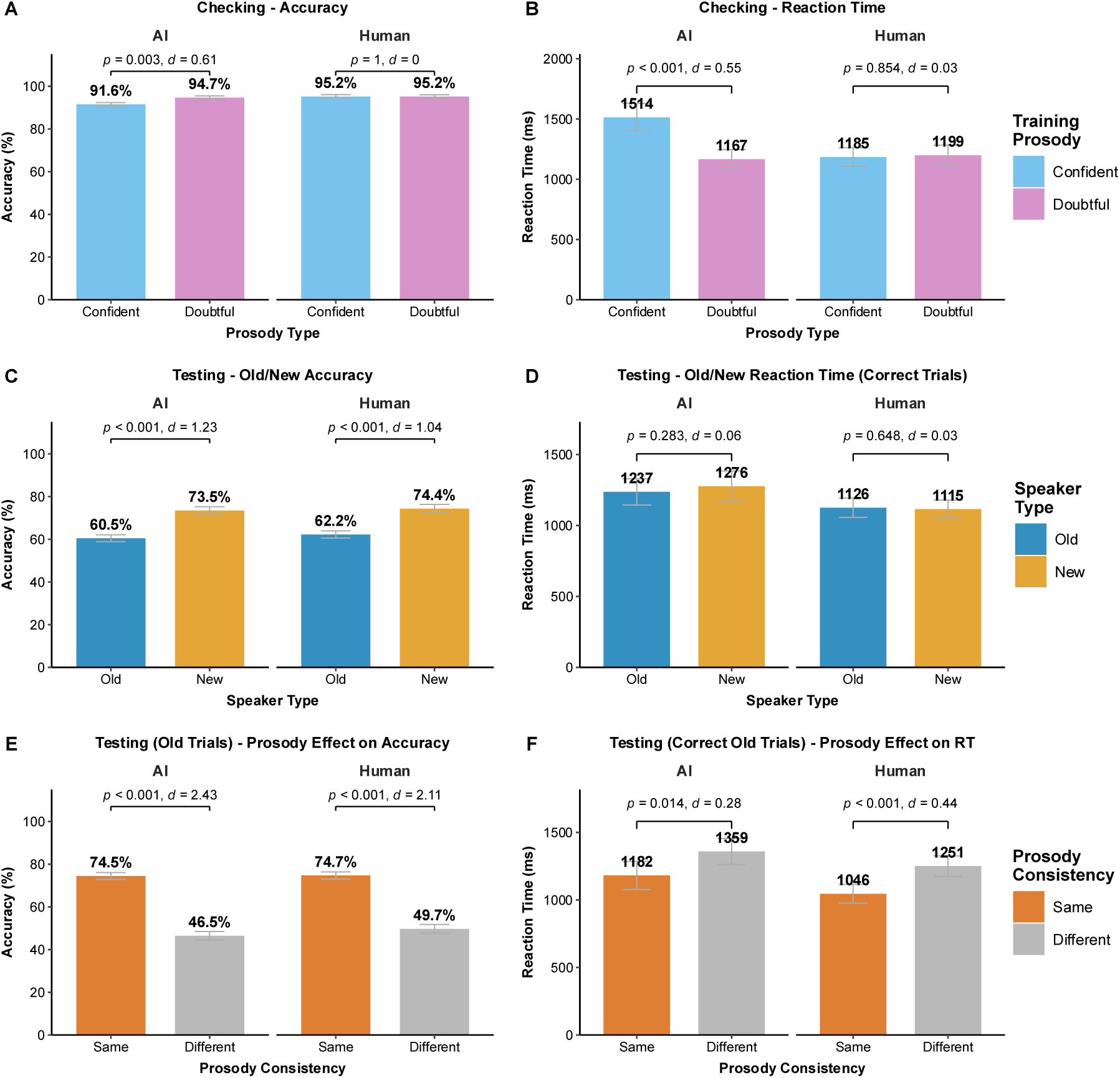
Behavioral performance in voice identity recognition. *Note.* **(A-B)** Accuracy and reaction time during the checking phase, where listeners identified speakers by selecting among three names. **(C-D)** Accuracy and reaction time (correctly old/new trials only) during the testing phase, where listeners judged whether voices belonged to old (trained) or new (untrained) speakers. **(E)** Accuracy for old speaker trials when prosody was the same vs. different between training and testing. (**F**) Reaction time for correctly recognized old speakers when prosody was the same vs. different. Error bars represent ±1 standard error of the mean.

In the Testing phase, participants were expected to classify the three trained familiar speakers as Old and the three untrained novel speakers as New, focusing only on speaker identity. Recognition accuracy was significantly lower for Old compared to New speakers in both human voices (old: 62.2%, new: 74.4%, *z* = –14.16, *p* < .001, *d* = 1.04) and AI voices (old: 60.5%, new: 73.5%, *z* = –15.15, *p* < .001, *d* = 1.23). Reaction times showed no difference between Old and New speakers for either voice type. See **Figure 3C-D**.

Listeners learned speaker identities in either confident or doubtful prosody. During testing, they encountered both confident and doubtful prosodies. Among Old speaker trials, those with the same prosody as training/checking yielded significantly higher accuracy than those with different prosody in both human voices (same: 74.7%, different: 49.7%, *z* = 20.61, *p* < .001, *d* = 2.11) and AI voices (same: 74.5%, different: 46.5%, *z* = 23.07, *p* < .001, *d* = 2.43). Similarly, reaction times (analyzed on correct trials only) were significantly faster for same vs. different prosody conditions in both human voices (same: 1046 ms, different: 1251 ms, *z* = –3.94, *p* < .001, *d* = 0.44) and AI voices (same: 1182 ms, different: 1359 ms, *z* = – 2.45, *p* = .014, *d* = 0.28). Notably, accuracy in the different prosody conditions dropped to chance level for both voice types. See **Figure 3E-F**.

### 3.2. Temporal MVPA results

For correctly judged trials, AI voices showed robust neural discrimination of old vs. new speakers in three significant clusters: 662-702 ms (AUC = 0.519 ± 0.052, *p* = .029), 758-844 ms (AUC = 0.525 ± 0.048, *p* = .006), and 866-1498 ms (AUC = 0.520 ± 0.050, *p* < .001), with peak decoding at 682 ms (AUC = 0.537). Human voices showed no significant discrimination (peak AUC = 0.527 at 1392 ms, all *p*s > .05; see **Figure 4A-B**, **E**). We also examined prosody consistency effects among correctly recognized old speakers (same vs. different prosody relative to training), but found no significant clusters for either voice type (all *p*s > .05), despite substantial behavioral differences (see **Figure 3E** and **Figure 4C-D**, **F**).

**Figure 4.**
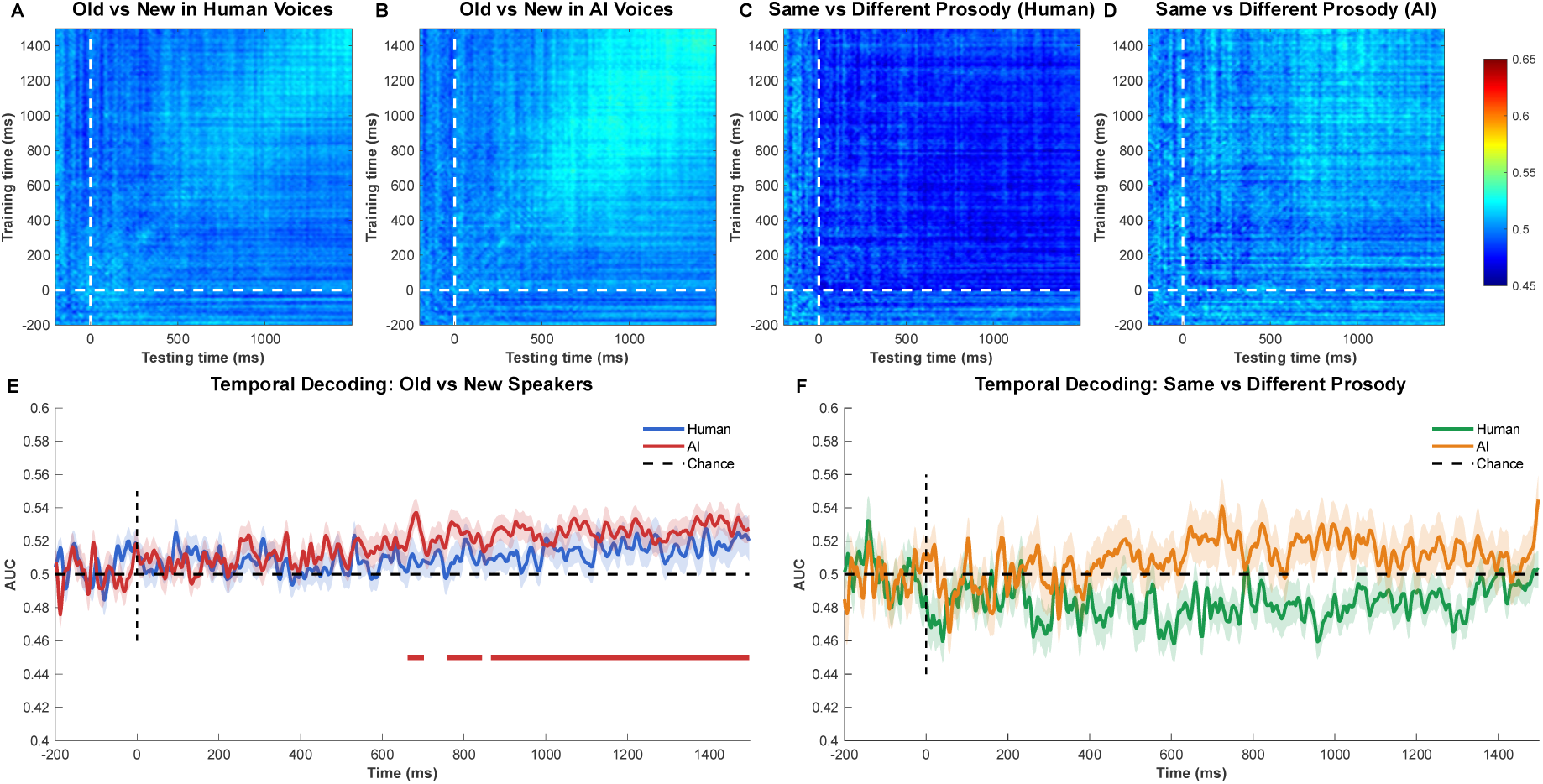
Time-resolved MVPA results for speaker recognition. *Note.* **(A-B)** Time generalization matrices (TGMs) for decoding old vs. new speakers (correct trials) in human and AI voices. **(C-D)** TGMs for decoding same vs. different prosody (relative to training) among correctly recognized old speakers. **(E)** Temporal decoding trajectories for old vs. new speakers; colored bars indicate significant clusters. **(F)** Temporal decoding trajectories for same vs. different prosody; no significant clusters identified. Dashed line = chance level (AUC = 0.50).

To identify electrodes contributing most to classifier discrimination, we applied Haufe transformation to the LDA weights and selected the top 10 contributing electrodes within each significant temporal window for AI voices. For the earlier windows (662-702 ms and 758-844 ms), contributing electrodes concentrated over posterior scalp regions, with Pz showing the strongest contribution. For the later window (866-1498 ms), electrodes spanned right frontal and posterior regions, with Pz remaining among the top contributors. These time windows and electrode selections were applied to human voices for parallel visualization and statistical analysis (**Figure 5**).

**Figure 5.**
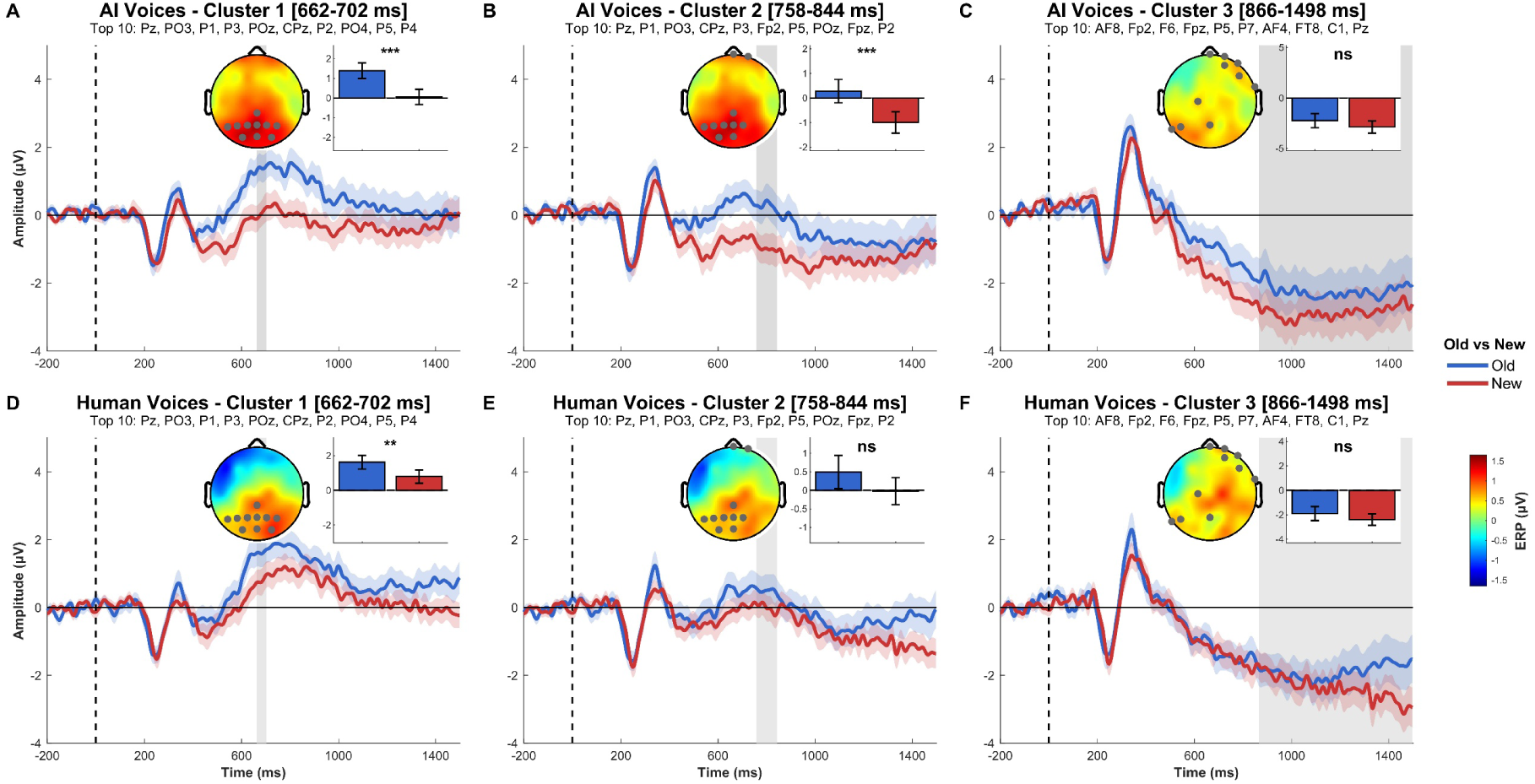
ERP amplitudes for top contributing electrodes in MVPA-identified windows. *Note.* **(A-C)** ERPs for AI voices in three significant temporal clusters. **(D-F)** ERPs for human voices using the same time windows and electrodes. Each panel shows ERP time course, topographic map of old-new difference, and mean amplitude comparison. Gray shading = analyzed window. \*\*\**p* < .001, \*\**p* < .01, \**p* < .05, ns = not significant.

To validate the neural distinctions identified by MVPA, we extracted mean ERP amplitudes within each significant temporal cluster and conducted linear mixed-effects regression comparing old vs. new speakers. For AI voices, old speakers elicited significantly more positive amplitudes than new speakers in the first two windows (cluster 1 [662-702 ms]: *F*(1, 7525) = 25.07, *p* < .001, β = 1.340; cluster 2 [758-844 ms]: *F*(1, 7540) = 16.38, *p* < .001, β = 1.250). For human voices, only the first window showed a significant old/new difference (cluster 1: *F*(1, 7589) = 10.05, *p* = .002, β = 0.828), while the second window showed a marginal trend (*F*(1, 7592) = 2.80, *p* = .094, β = 0.489). The later window (866-1498 ms) showed no significant old/new difference for either voice type (AI: *F*(1, 7537) = 2.40, *p* = .122, β = 0.626; Human: *F*(1, 7588) = 1.79, *p* = .181, β = 0.470). Also see **Tables S1** and **S2** for statistical details.

### 3.3. Speech-content-independent Old vs. New effect (N250/P300/LPC)

We analyzed mean amplitudes at nine scalp electrodes (F3, Fz, F4, C3, Cz, C4, P3, Pz, P4) across three time windows (N250: 200-280 ms; P300: 300-380 ms; LPC: 400-800 ms) using linear mixed-effects regression (see **Table S3**, **Table S4**, **Figure 7**, **Figure 6**, and **Figure 8**).

**Figure 6.**
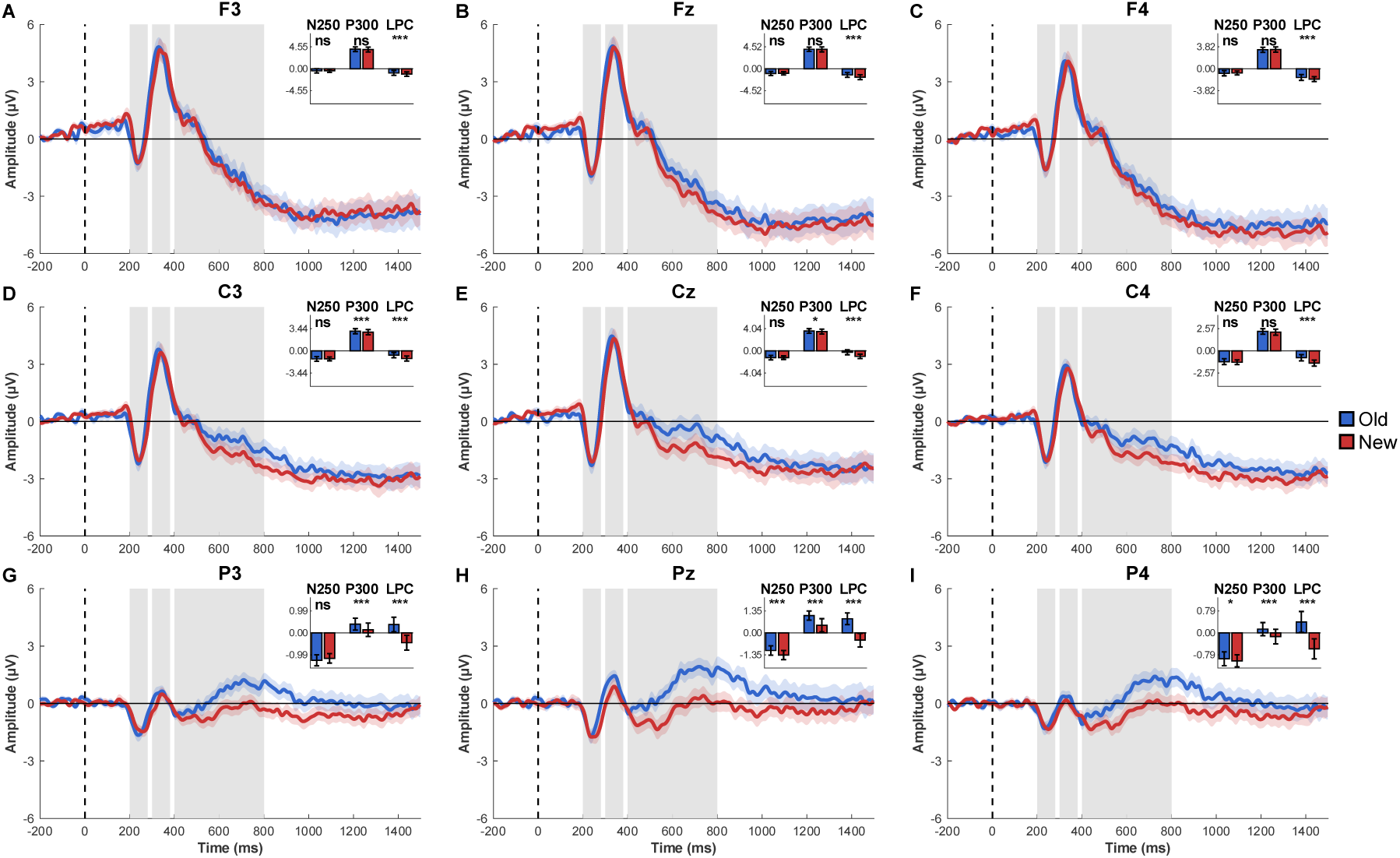
ERP waveforms for old vs. new speaker recognition in AI voices. *Note.* **(A-I)** ERPs at nine scalp electrodes. Gray shading indicates analyzed time windows (N250: 200-280 ms; P300: 300-380 ms; LPC: 400-800 ms). Inset bar plots show mean amplitudes for each window. Significance markers: \*\*\**p* < .001, \*\**p* < .01, \**p* < .05, ns = not significant.

**Figure 7.**
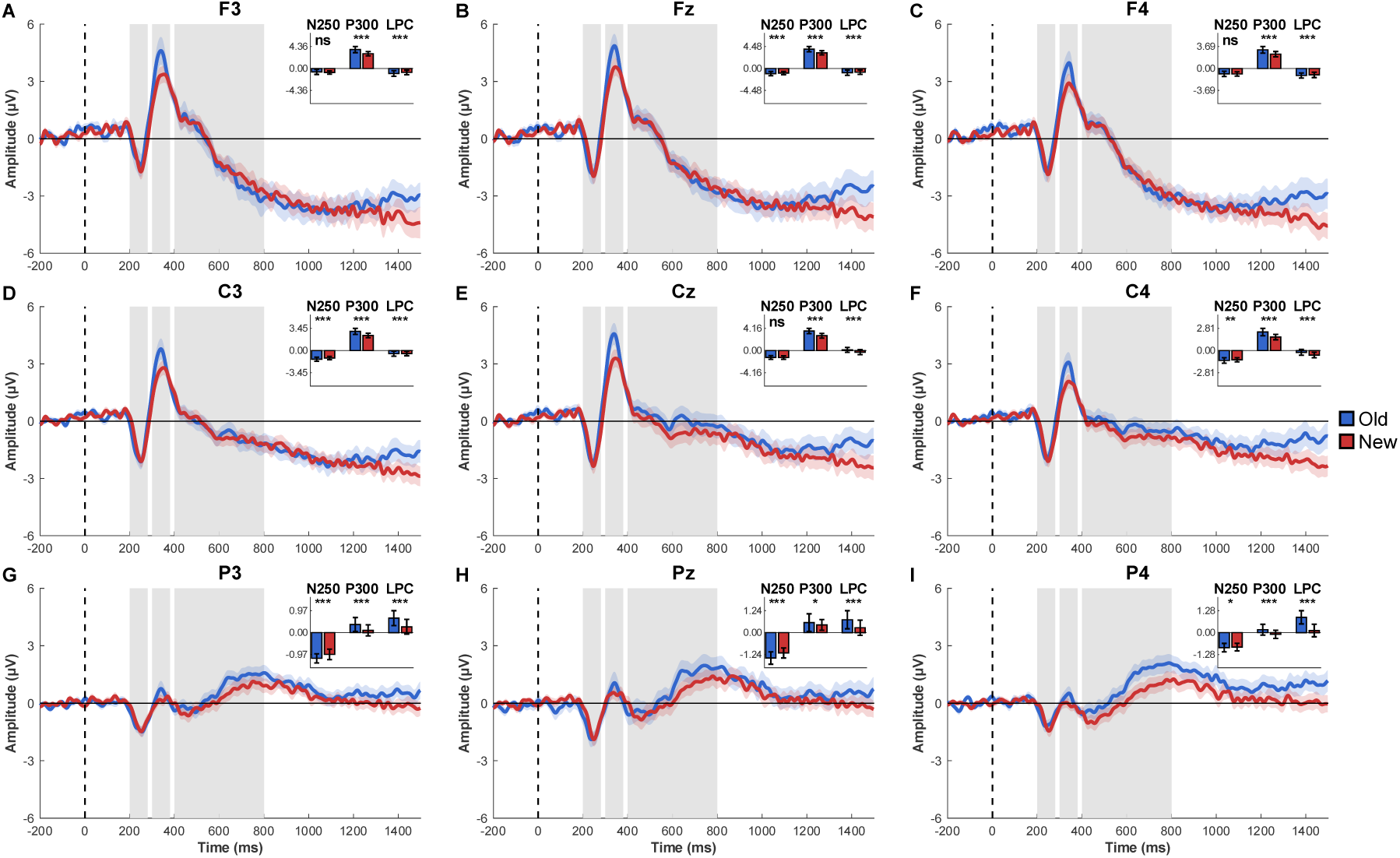
ERP waveforms for old vs. new speaker recognition in human voices. *Note.* **(A-I)** ERPs at nine scalp electrodes. Gray shading indicates analyzed time windows (N250: 200-280 ms; P300: 300-380 ms; LPC: 400-800 ms). Inset bar plots show mean amplitudes for each window. Significance markers: \*\*\**p* < .001, \*\**p* < .01, \**p* < .05, ns = not significant.

**Figure 8.**
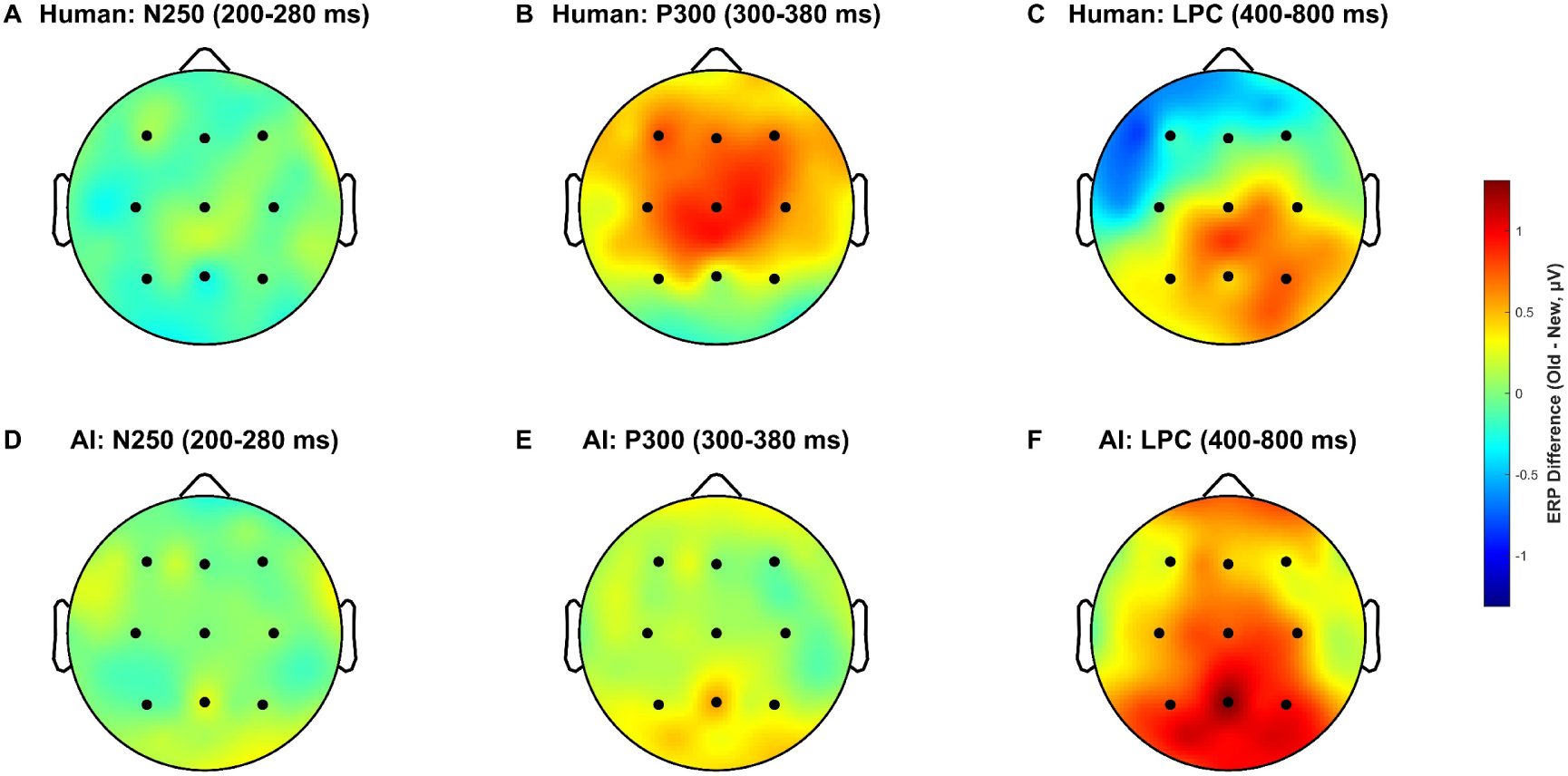
ERP difference maps for old-new speaker recognition. *Note.* (**A**-**C**) Human voices in three time windows (N250, P300, LPC). (**D**-**F**) AI voices in the same windows. Black circles mark the nine analyzed electrodes.

In the N250 window, AI voices showed that old speakers elicited significantly greater negativity than new speakers at Pz (*F*(1, 309218) = 35.82, β = 0.29, *p* < .001) and P4 (*F*(1, 308515) = 4.52, β = 0.08, *p* = .033), with no significant differences at other electrodes. For human voices, old speakers elicited significantly greater negativity at six electrodes (ordered by effect size): Pz (*F*(1, 311628) = 48.93, β = –0.32, *p* < .001), P3 (*F*(1, 311890) = 40.70, β = –0.24, *p* < .001), C3 (*F*(1, 311860) = 28.83, β = –0.22, *p* < .001), Fz (*F*(1, 311921) = 12.38, β = –0.18, *p* < .001), C4 (*F*(1, 311938) = 8.58, β = –0.12, *p* = .003), and P4 (*F*(1, 311738) = 6.24, β = –0.09, *p* = .013), with no significant effects at Cz, F3, and F4.

In the P300 window, AI voices showed that old speakers elicited significantly greater positivity than new speakers at five electrodes (ordered by effect size): Pz (*F*(1, 309411) = 137.83, β = 0.59, *p* < .001), P3 (*F*(1, 308966) = 47.15, β = 0.28, *p* < .001), P4 (*F*(1, 309072) = 45.46, β = 0.28, *p* < .001), C3 (*F*(1, 309130) = 12.06, β = 0.16, *p* < .001), and Cz (*F*(1, 309257) = 6.37, β = 0.13, *p* = .012), with no significant differences at frontal electrodes. For human voices, old speakers elicited significantly greater positivity at all nine electrodes (ordered by effect size): Cz (*F*(1, 311951) = 294.46, β = 0.83, *p* < .001), C4 (*F*(1, 311904) = 193.24, β = 0.60, *p* < .001), C3 (*F*(1, 311896) = 163.91, β = 0.57, *p* < .001), F4 (*F*(1, 311918) = 160.54, β = 0.66, *p* < .001), Fz (*F*(1, 311930) = 136.27, β = 0.64, *p* < .001), F3 (*F*(1, 311837) = 134.07, β = 0.69, *p* < .001), P4 (*F*(1, 311780) = 32.22, β = 0.22, *p* < .001), P3 (*F*(1, 311890) = 22.68, β = 0.19, *p* < .001), and Pz (*F*(1, 311777) = 4.36, β = 0.10, *p* = .037).

In the LPC window, AI voices showed that old speakers elicited significantly greater positivity than new speakers at all nine electrodes (ordered by effect size): Pz (*F*(1, 1518133) = 2715.60, β = 1.35, *p* < .001), P4 (*F*(1, 1517857) = 2364.50, β = 1.02, *p* < .001), P3 (*F*(1, 1517390) = 1761.10, β = 0.88, *p* < .001), Cz (*F*(1, 1517675) = 955.48, β = 0.78, *p* < .001), C4 (*F*(1, 1517723) = 867.51, β = 0.66, *p* < .001), C3 (*F*(1, 1517294) = 548.98, β = 0.54, *p* < .001), Fz (*F*(1, 1517963) = 247.42, β = 0.44, *p* < .001), F4 (*F*(1, 1517178) = 122.73, β = 0.30, *p* < .001), and F3 (*F*(1, 1516924) = 38.85, β = 0.19, *p* < .001). For human voices, old speakers elicited significantly greater positivity at five posterior and central electrodes (ordered by effect size): P4 (*F*(1, 1529408) = 1378.20, β = 0.74, *p* < .001), Pz (*F*(1, 1529319) = 370.31, β = 0.48, *p* < .001), P3 (*F*(1, 1529579) = 303.10, β = 0.35, *p* < .001), Cz (*F*(1, 1529560) = 272.36, β = 0.40, *p* < .001), and C4 (*F*(1, 1529518) = 183.63, β = 0.29, *p* < .001). Notably, at frontal and left-central electrodes, old speakers elicited significantly greater negativity: F3 (*F*(1, 1529316) = 98.50, β = –0.31, *p* < .001), Fz (*F*(1, 1529548) = 44.93, β = – 0.18, *p* < .001), F4 (*F*(1, 1529480) = 40.18, β = –0.17, *p* < .001), and C3 (*F*(1, 1529507) = 14.13, β = –0.09, *p* < .001).

To examine effect size progression across time windows, we calculated Spearman rank correlations between time window order (N250, P300, LPC) and ω² values across all nine electrodes. AI voices showed a significant positive correlation (ρ = 0.735, *p* < .001), indicating progressive strengthening from N250 (mean ω² = 0.000015) through P300 (0.000088) to LPC (0.000702), with significant electrodes expanding from 2 to 9. In contrast, human voices showed no significant linear trend (ρ = 0.321, *p* = .103), with effect sizes peaking at P300 (0.000403) before declining at LPC (0.000196), despite maintaining significance at all nine electrodes from P300 onward.

To summarize, four key patterns emerged. First, both voice types demonstrated reliable old/new discrimination across all three time windows, with convergent parietal engagement (Pz) in the LPC window. Second, spatial organization differed markedly: AI voices maintained consistent posterior-centered patterns with uniform positive polarity, while human voices showed more widespread distributions across scalp regions and late frontal polarity reversal. Third, AI voices showed larger neural responses than human voices, with progressive electrode recruitment (N250: 2 electrodes; P300: 5 electrodes; LPC: 9 electrodes), whereas human voices showed broader early engagement that stabilized (N250: 6 electrodes; P300-LPC: 9 electrodes). Fourth, temporal dynamics diverged: AI voices showed significant effect size increases across windows (ρ = 0.735, *p* < .001), reflecting progressive strengthening of neural discrimination, while human voices showed no significant linear trend (ρ = 0.321, *p* = .103), with early robust engagement.

### 3.4. Speaker-specific prosody expectation effect (late time window)

We analyzed mean amplitudes at nine scalp electrodes (F3, Fz, F4, C3, Cz, C4, P3, Pz, P4) within the late window (500-900 ms) using linear mixed-effects regression to examine prosody consistency effects (same vs. different prosody relative to training). See **Table S5**, **Table S6**, **Figure 10**, **Figure 9**, and **Figure 11**.

**Figure 9.**
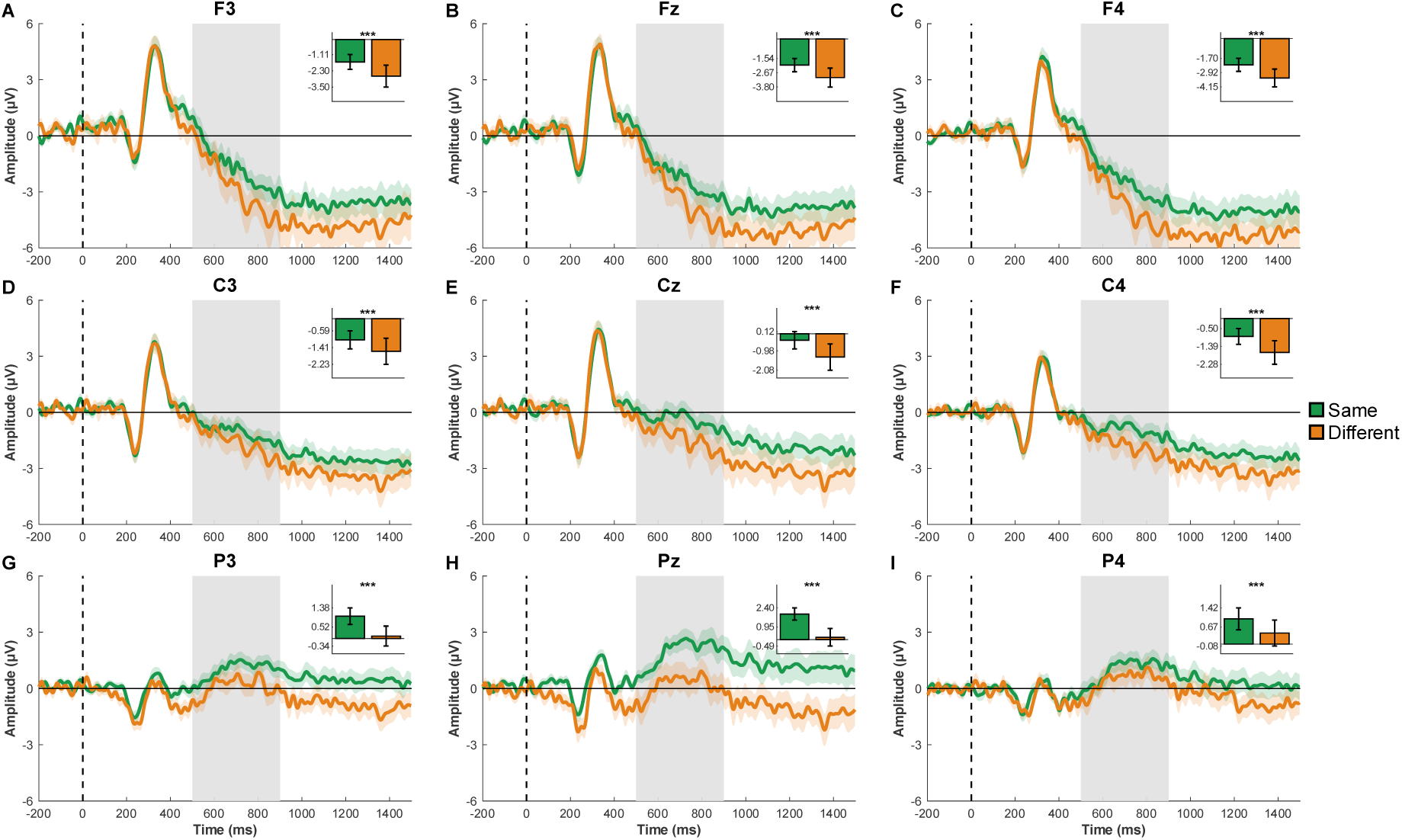
ERP waveforms for speaker-specific prosody effects in AI voices. *Note.* **(A-I)** ERPs at nine scalp electrodes comparing same vs. different prosody relative to training. Gray shading indicates the late window (500-900 ms) where prosody violations elicit neural responses. Inset bar plots show mean amplitudes. \*\*\**p* < .001, \*\**p* < .01, \**p* < .05, ns = not significant.

**Figure 10.**
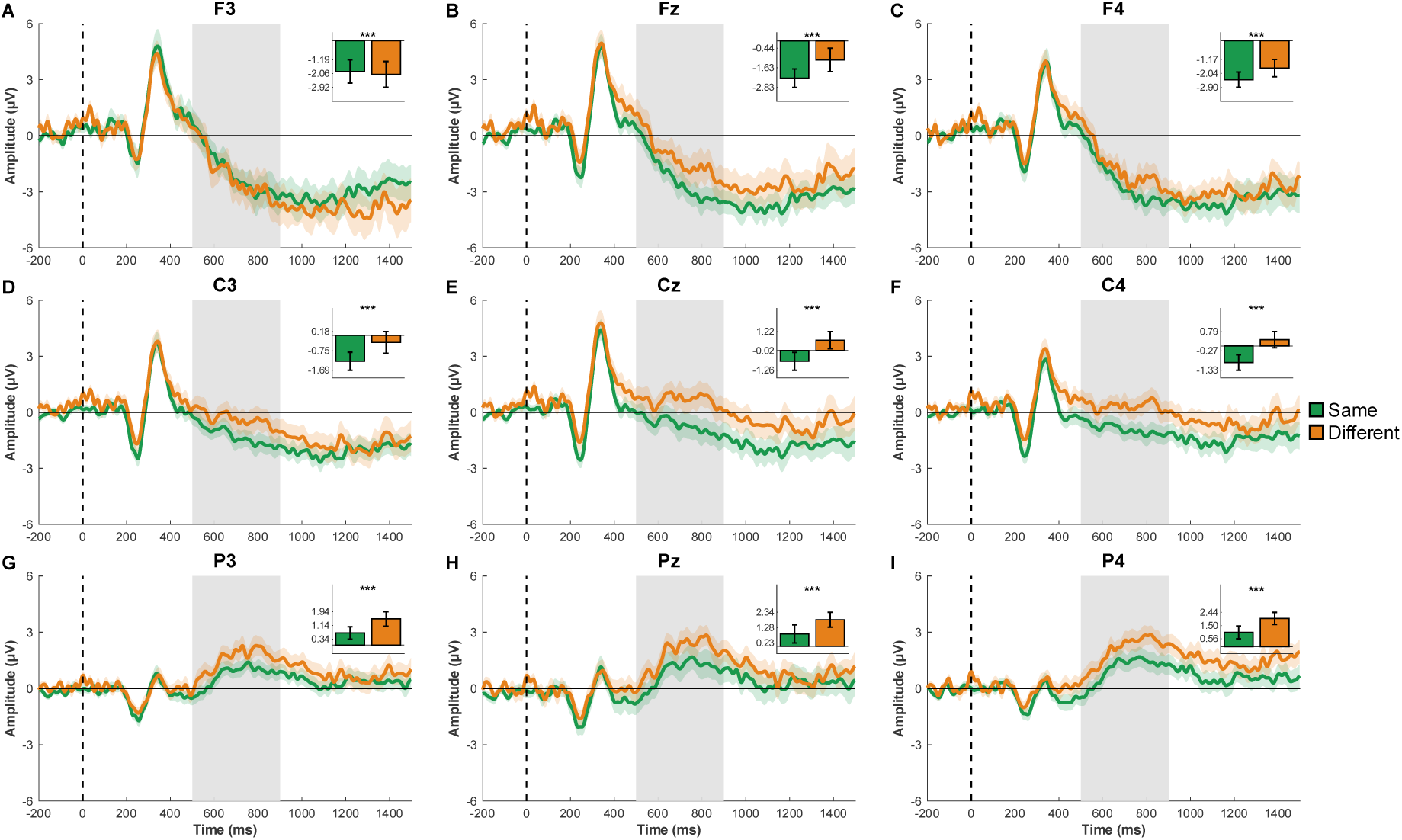
ERP waveforms for speaker-specific prosody effects in human voices. *Note.* **(A-I)** ERPs at nine scalp electrodes comparing same vs. different prosody relative to training. Gray shading indicates the late window (500-900 ms) where prosody violations elicit neural responses. Inset bar plots show mean amplitudes. \*\*\**p* < .001, \*\**p* < .01, \**p* < .05, ns = not significant.

**Figure 11.**
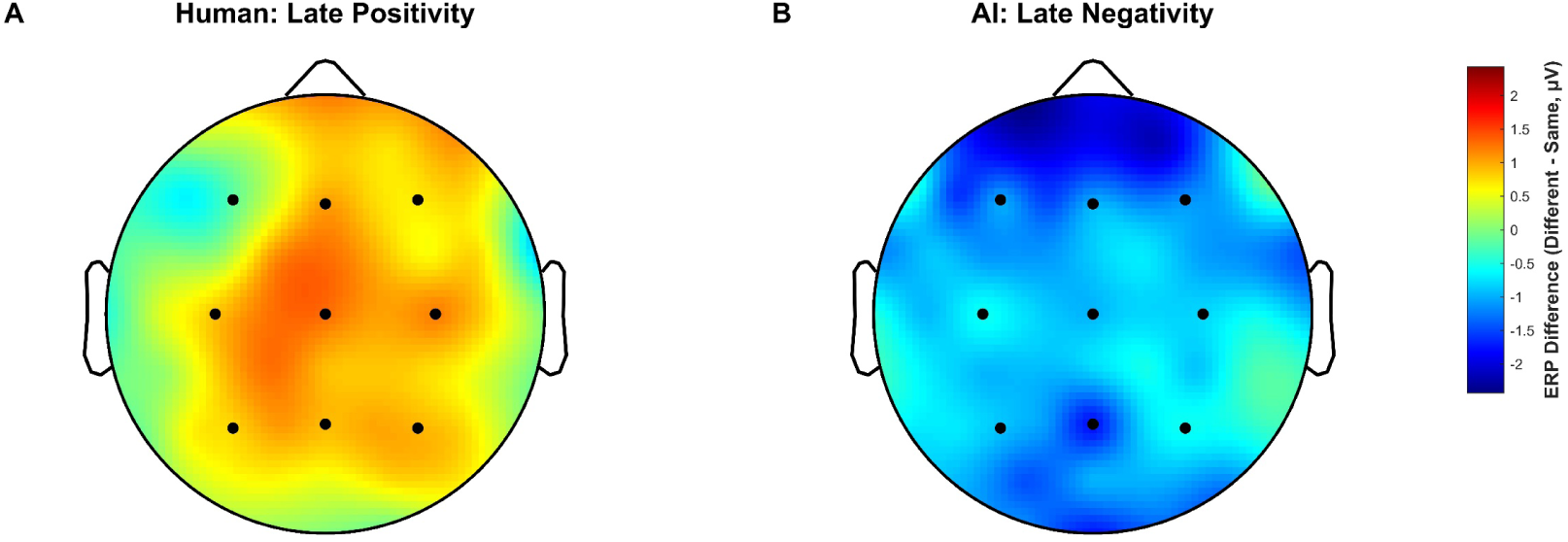
ERP difference maps for speaker-specific prosodic expectancy violations in correctly recognized old speakers. *Note.* **(A)** Human voices show late positivity (500-900 ms). **(B)** AI voices show late negativity (500-900 ms). Black dots indicate the nine analyzed electrodes (Fz, FCz, Cz, CPz, Pz, P3, P4, POz, Oz).

In the late window, AI voices showed that same prosody elicited significantly greater negativity than different prosody at all nine electrodes (ordered by effect size): Pz (*F*(1, 684902) = 910.16, β = 1.20, *p* < .001), P3 (*F*(1, 684486) = 415.27, β = 0.67, *p* < .001), F4 (*F*(1, 682677) = 196.42, β = 0.59, *p* < .001), C4 (*F*(1, 683909) = 152.71, β = 0.43, *p* < .001), P4 (*F*(1, 684568) = 130.58, β = 0.38, *p* < .001), Cz (*F*(1, 684303) = 114.86, β = 0.42, *p* < .001), Fz (*F*(1, 684344) = 98.06, β = 0.43, *p* < .001), F3 (*F*(1, 682273) = 95.36, β = 0.45, *p* < .001), and C3 (*F*(1, 684377) = 23.33, β = 0.18, *p* < .001).

For human voices, same prosody elicited significantly greater positivity than different prosody at eight electrodes (ordered by effect size): C4 (*F*(1, 697828) = 1252.50, β = –1.17, *p* < .001), P4 (*F*(1, 697779) = 934.66, β = –0.93, *p* < .001), Cz (*F*(1, 697826) = 816.96, β = – 1.07, *p* < .001), P3 (*F*(1, 697844) = 527.43, β = –0.70, *p* < .001), Fz (*F*(1, 697747) = 434.09, β = –0.89, *p* < .001), C3 (*F*(1, 697823) = 428.29, β = –0.72, *p* < .001), Pz (*F*(1, 697838) = 385.43, β = –0.74, *p* < .001), and F4 (*F*(1, 697703) = 375.42, β = –0.79, *p* < .001). Notably, F3 showed an opposite pattern, with same prosody eliciting greater negativity (*F*(1, 697694) = 18.45, β = 0.22, *p* < .001).

Overall, both voice types showed robust prosody consistency effects across all nine electrodes, but with opposite polarities: AI voices showed late negativity for prosody violations, while human voices showed late positivity.

## 4. Discussion

Our findings provide mixed support for the three hypotheses. Hypothesis 1 (speech-content-independent recognition) was partially supported: robust parietal old/new effects emerged even when sentences changed completely between training and testing, confirming abstract identity representations beyond episodic memory. Although we anticipated replicating speech-independent beta band oscillations (16-17 Hz), time-frequency MVPA revealed no significant clusters for either voice type. Hypothesis 2 (speaker-specific prosody expectations) was also supported: prosodic violations elicited robust late components, confirming that listeners bind prosodic patterns to speaker identities. Hypothesis 3 (AI-human comparability) received mixed support: both voice types showed comparable parietal old/new effects behaviorally, yet MVPA revealed stronger old/new discrimination for AI voices than human voices. Moreover, while both showed late components for prosodic violations, the opposite polarities (human positivity vs. AI negativity) contradicted our expectation of comparable mechanisms, instead revealing fundamentally distinct processing routes.

### 4.1. Content-independent speaker recognition and the contribution of name-labeling to identity encoding

In previous old/new studies, <u>Zäske et al. (2014)</u> reported ∼62-70% accuracy (depending on block half) using neutral speech without prosodic variation, while <u>Xu and Armony (2021)</u> found accuracy dropping sharply toward chance (∼39-49%) when prosody differed between training and testing (neutral vs. fearful). Our study used naturally varying prosodic styles (confident vs. doubtful), yet replicated the accuracy pattern of <u>Zäske et al. (2014)</u> and substantially improved upon <u>Xu and Armony (2021)</u>.

Having established above-chance behavioral performance with robust identity encoding, we analyzed EEG data from correctly judged old/new trials to investigate the neural mechanisms underlying content-independent voice recognition. Multiple converging lines of evidence pointed to Pz and parietal regions as central sites for identity processing. First, MVPA identified Pz as a major contributor in the late decoding window for both voice types. Second, univariate analyses revealed significant old/new effects at Pz. Third, effect-size rankings consistently placed Pz among the strongest responders across time windows. Critically, unlike <u>Zäske et al. (2014)</u>, who observed parietal LPC only when training and testing used identical sentences, our study demonstrated robust parietal LPC effects even when speech content changed completely between training and testing. This finding indicates that the parietal LPC can reflect speech-content-independent identity retrieval rather than episodic memory for specific utterances.

We speculate that these behavioral and neural differences arise from how identity was encoded in our paradigm. In previous studies using passive exposure (<u>Xu & Armony, 2021</u>; <u>Zäske et al., 2017</u>; <u>Zäske et al., 2014</u>), listeners heard voices repeatedly without explicit identity labeling, resulting in near-chance behavioral accuracy and limited neural discrimination when content varied. In contrast, our listeners learned each voice together with a name and we ensured this name-voice association was fully established through checking verification (<u>Lavan, Knight, et al., 2019</u>). This explicit name-labeling procedure elevated both behavioral recognition accuracy and neural discriminability.

As such, we speculate that names may play a more direct role than other labels in facilitating identity consolidation. Unlike occupations or semantic attributes, names carry no inherent meaning and cannot be retrieved directly from perceptual cues such as faces or voices; instead, they must be accessed through person-identity codes (<u>McWeeny et al., 1987</u>). Because names are cognitively unique identity labels that require stronger encoding than other person information, successfully binding a name to a newly learned voice may strengthen person-level representations. In the framework of the looping mechanism (<u>Maguinness et al., 2018</u>), where repeated recognition episodes progressively strengthen voice prototype representations, name-based learning (<u>McWeeny et al., 1987</u>) may establish more robust identity anchors than arbitrary symbols or numbers, facilitating the consolidation of stable voice identities. However, this speculation lacks direct supporting evidence and requires controlled comparisons in future research.

So far, we cannot definitively attribute these neural signatures solely to name-labeling. On one hand, our trained-familiar voices with name associations still differ qualitatively from genuinely intimate voices. Voices of close family members, long-term friends (<u>Plante-Hébert et al., 2021</u>), or well-known public figures (<u>Rinke et al., 2022</u>) carry autobiographical, affective, and deeply consolidated semantic associations that likely exceed what can be induced through brief laboratory training. On the other hand, our name-labeling procedure may have achieved its effects through mechanisms shared with other explicit identity anchors. Previous studies have linked voices to icons (<u>Cooper et al., 2024</u>; <u>Perrachione et al., 2011</u>) or numbers (<u>Xie & Myers, 2015</u>), and these alternative labels also enhance familiarity relative to passive exposure. Whether names confer unique advantages over other explicit labels, or whether any salient identity anchor would produce similar content-independent representations, remains an empirical question for future research. Controlled comparisons directly contrasting name-labeling with icon-labeling or number-labeling within the same paradigm would be necessary to isolate the specific contribution of names.

Nevertheless, our LPC findings suggest a possible pathway through which name-labeling may elevate newly learned voices toward intimate familiarity. In <u>Plante-Hébert et al. (2021)</u>, voice familiarity spans three levels: unfamiliar voices; trained-to-familiar voices acquired through repeated exposure without explicit identity labeling; and intimately familiar voices such as family and friends. Intimate familiarity produces two key neural signatures: strongly parietal spatial distributions extending from Pz across posterior midline electrodes, and robust late positive components reflecting retrieval of rich person-specific representations. Our old/new effects exhibited similar patterns. The spatial distribution was strongly parietal, with Pz as the strongest contributor and additional posterior sites showing reliable discrimination. Moreover, we observed a robust LPC even when speech content varied, suggesting listeners accessed stable voice identity representations rather than episodic traces of specific sentences. These convergences raise the possibility that pairing a voice with a name and ensuring listeners correctly learn this association elevates newly learned voices toward a level of familiarity that approximates key aspects of intimate voice processing. Thus, we speculate that name-voice association learning could represent a foundational process by which new acquaintances transition from strangers to socially familiar individuals (<u>Lavan & McGettigan, 2023</u>; <u>Maguinness et al., 2018</u>; <u>Sidtis & Kreiman, 2012</u>).

We also note that <u>Zäske et al. (2014)</u> additionally reported beta band (16-17 Hz) effects during 290-370 ms for old vs. new voices using univariate power analyses at specific electrode clusters. Our time-frequency MVPA (**Supplementary Analysis 1**) did not replicate this effect. Two methodological differences may account for this discrepancy. First, <u>Zäske et al. (2014)</u> employed univariate analyses testing frequency-specific power differences at individual electrodes, whereas our multivariate decoding classified conditions based on distributed spatial patterns across all electrodes, which may be less sensitive to localized frequency-specific effects. Second, our paradigm introduced prosodic variability (confident vs. doubtful) that was absent in the neutral-prosody-only design of <u>Zäske et al. (2014)</u>. Future work could test the replicability of these beta effects.

### 4.2. Speaker-specific identity representations remain stable across prosodic variation

In predictive processing, the brain continuously generates predictions to minimize mismatches between expected and received sensory inputs (<u>Friston & Kiebel, 2009</u>). We analyzed ERP data from trials where participants successfully identified old speakers and found robust late components, supporting our hypothesis that prosody serves as a speaker-specific predictive cue. Specifically, old speaker-based prosodic violations elicited robust late neural responses in both voice types, but with opposite polarities: human voices showed enhanced late positivity, while AI voices showed enhanced late negativity.

For human voices, this late positivity reflects pragmatic repair mechanisms similar to those observed when listeners reconcile conflicting vocal confidence cues (<u>Jiang & Pell, 2016</u>). When prosodic violations remain communicatively coherent, as in our paradigm where speaker identity stays constant despite prosodic shifts, listeners engage inferential reinterpretation rather than rejecting the speaker representation. This pattern aligns with P600 components elicited by violations of speaker-specific communicative styles, including syntactic structure preferences (<u>Kroczek & Gunter, 2021</u>), ironic tendencies (<u>Regel et al., 2010</u>), and gesture patterns (<u>Obermeier et al., 2015</u>). Across these domains, late positivities emerge when violations are recoverable, that is, when the mismatch can be reconciled with stored speaker representations without rejecting the speaker’s identity or communicative intent. Our findings extend this mechanism to prosodic style, demonstrating that listeners bind prosodic patterns to speaker identities and flexibly update expectations when encountering within-speaker prosodic variation.

In contrast, AI voices elicited late negativity, reflecting effortful reprocessing mechanisms. This pattern parallels findings that atypical vocal signals create processing challenges. <u>Jiang et al. (2020)</u> found that compared to local Canadian English, non-native regional accents (Australian English) and L2-accented speech (French-accented English) combined with doubtful vocal confidence elicited sustained late negativity, reflecting integration difficulties when constructing speaker representations from atypical vocal patterns. Similarly, AI-generated voices may be processed as a distinct type of speech signal analogous to unfamiliar accents (<u>Roswandowitz et al., 2024</u>): although listeners successfully recognize speaker identity, the synthetic origin creates a baseline processing challenge. When prosodic violations occur within this already-atypical vocal framework, listeners must reconcile expectations across prosodic styles while also processing artificially-generated acoustic patterns. This dual challenge pushes processing toward effortful reinterpretation routes involving inhibition and recomputation (<u>Jiang et al., 2013</u>), rather than the flexible updating mechanisms available for natural human voices where only prosodic expectations require adjustment.

The significance of these findings becomes evident when considering our cloning procedure. Human prosodic variations originated from the same speaker across identical sentences, maintaining consistent identity. AI voices presented a more stringent test: each speaker’s confident and doubtful clones were independently trained from separate 15-sentence corpora, with no cross-training between prosodic styles. The AI models then generated 123 novel sentences they had never encountered during training, which human speakers subsequently recorded one month later. Despite this algorithmic separation and the generative nature of AI speech, both voice types elicited robust late-window ERP differences for prosodic violations, demonstrating that listeners formed unified identity representations. This provides direct evidence that speaker identity remains sufficiently stable across prosodic variations, provided the variations are not overly expressive (<u>Lavan, Burston, et al., 2019</u>). Both AI generative models and human listeners successfully extracted shared identity characteristics despite within-speaker prosodic variation. These findings support theories of flexible within-speaker identity processing (<u>Lavan, Burton, et al., 2019</u>).

Our study extends research on speaker-specific communicative styles by examining prosody, which differs critically from syntactic or other stylistic cues in that prosody additionally shifts speaker identity representations through VTL and F0 adjustments (<u>Lavan, Knight, et al., 2019</u>). This creates a unique processing challenge: listeners must match the current speaker’s identity representation with stored representations of familiar talkers despite prosodic differences (<u>Lavan & McGettigan, 2023</u>), which requires greater effort than matching identical prosodic styles (e.g., both neutral). Only after successful identity retrieval can listeners process pragmatic information conveyed by prosodic violations. Thus, prosody introduces an interaction between identity-level adaptation to within-speaker acoustic variation and pragmatic-level processing of communicative style, demonstrating the flexible capacity of the auditory system to accommodate both levels simultaneously.

### 4.3. AI voices as methodological tools for isolating identity processing?

Despite similar behavioral patterns for the old/new effects, our MVPA analyses revealed a notable pattern: AI voices showed three significant decoding clusters for old/new discrimination, whereas human voices showed no significant clusters independently (**Figure 5**). However, human voices showed comparable old/new ERP effects when analyzed using AI-identified time windows and electrodes in linear mixed-effects models. This asymmetry likely stems from reduced prosodic variability in AI voices. Although both voice types conveyed prosodic distinctions (see <u>Chen et al. (2025)</u> and **Figure 1**), human voices exhibited substantially larger prosodic differences. Because our old/new analyses collapsed across prosodic conditions, greater prosodic heterogeneity in human voices may have obscured identity patterns in MVPA, whereas AI voices provided cleaner signals for identity extraction.

Beyond prosodic variation, AI voices may exhibit greater internal homogeneity than human voices (<u>Chen et al., 2024</u>). Natural human speech carries rich variation in paralinguistic, emotional, and social-affective dimensions that engage broader neural networks (<u>Roswandowitz et al., 2024</u>).While this richness reflects the complexity of human voice processing, it can introduce confounds when investigating specific cognitive processes. AI-generated voices, by minimizing these extraneous factors, may serve as valuable methodological tools for isolating core identity processing mechanisms, such as investigating neural sensitivity to different levels of voice familiarity (<u>Ma et al., 2026</u>; <u>McGettigan et al., 2025</u>) without confounding effects from emotional, prosodic, or social-affective variation.

Our prosody consistency effect further hints at this methodological advantage. Although neither voice type showed significant MVPA clusters for same vs. different prosody discrimination, AI voices maintained above-chance decoding accuracy while human voices fluctuated around chance levels. Human voices carry richer incidental variation such as pronunciation inconsistencies or accent features that introduce noise beyond the intended prosodic manipulation (<u>Jiang et al., 2020</u>; <u>Jiang et al., 2018</u>; <u>Leemann et al., 2018</u>; <u>Ulbrich & Mennen, 2016</u>). AI-generated voices, by delivering standardized outputs, maximize signal-to-noise ratio for experimentally manipulated dimensions. For example, <u>Di Cesare et al. (2022)</u> introduced a robotic voice lacking prosodic intonation as a control condition to isolate the contribution of human vocal prosody. AI voice synthesis similarly offers a useful tool for investigating how specific acoustic dimensions influence identity processing by isolating target variables while holding other factors constant.

Meanwhile, we need to highlight that univariate ERP analyses revealed early processing advantages for human voices that MVPA could not detect. In the N250 window (200-280 ms), human voices showed old/new discrimination across six electrodes vs. only two for AI voices, suggesting faster identity extraction. Conversely, MVPA identified significant decoding clusters for AI voices but not human voices. Thus, the two voice types engage temporally distinct processing trajectories: human voices demonstrate early robust discrimination, whereas AI voices show late-stage multivariate decoding advantages.

We also note that human voices demonstrated overall learning advantages in the Checking phase. Within AI voices, confident prosody yielded lower accuracy than doubtful prosody, whereas human voices showed no prosodic differences. This suggests AI confident voices have greater internal similarity among learned speakers despite our height-matched controls. These findings indicate identity learning operates holistically and is influenced by prosody, with future research needing to attend to differential prosodic effects when using AI voices.

Overall, while AI voices demonstrated MVPA advantages and human voices showed early univariate discrimination, both approaches converged on identifying Pz as a critical contributor. This suggests that despite temporally distinct trajectories, both voice types engage overlapping parietal substrates for identity processing. The choice should depend on research goals: AI voices for minimizing confounding variation, human voices for ecological validity and early processing dynamics.

### 4.4. Limitations and future directions

Our fixed block order (human blocks 1-4, then AI blocks 5-8) requires special attention, as earlier blocks typically show stronger identity encoding advantages than later blocks (<u>Xu & Armony, 2021</u>; <u>Zäske et al., 2014</u>), suggesting alternating human/AI blocks might be preferable. However, our Checking phase results (**Figure 3A**) justify this design. Despite appearing in later blocks with potential cumulative learning advantages, AI voices showed systematically lower accuracy than human voices, particularly for AI confident prosody. This persistent disadvantage despite later positioning suggests fundamental encoding differences. As discussed above, AI confident voices likely exhibited greater internal perceptual similarity among the three learned speakers. Had we alternated human/AI blocks while counterbalancing prosody order, AI voices might have shown even greater disadvantages, and this would have disrupted our human voice recognition baseline needed for comparison with prior exclusively-human-voice literature. As we examined only two pragmatically-marked prosodic types (confident and doubtful) rather than more basic emotional prosody such as happy or sad (<u>Larrouy-Maestri et al., 2025</u>), future research could also extend or replicate our paradigm using emotional prosody (<u>Xu & Armony, 2021</u>), while ensuring within-speaker variation remains within appropriate ranges (<u>Lavan, Burston, et al., 2019</u>).

Recall as well that our study did not directly compare neural responses between human and AI voices but instead examined parallel patterns within each voice type (old/new effects and speaker-specific prosodic expectation violations) to infer shared underlying mechanisms. As such, our current design cannot address several important questions: can listeners rapidly perceive the distinction between human and AI voices at early processing stages (<u>Jiang & Pell, 2024</u>; <u>Lavan et al., 2024</u>)? Does this human/AI discrimination depend on prosodic cues (<u>Kühne et al., 2020</u>; <u>Rodero & Lucas, 2023</u>)? Future research should investigate these questions to clarify the temporal dynamics and cognitive mechanisms underlying human vs. AI voice discrimination.

## 5. Conclusion

Our study addresses current gaps in voice processing research. First, prior work reported parietal old/new effects for voice recognition only when sentences were repeated at test (<u>Zäske et al., 2014</u>); we demonstrate these effects persist across different speech content, indicating abstract identity representations beyond episodic memory. Second, listeners construct predictive models of interlocutors’ communicative behaviors based on available social cues, forming expectations for speaker-specific patterns (<u>Kroczek & Gunter, 2021</u>; <u>Kroczek et al., 2019</u>; <u>Obermeier et al., 2015</u>; <u>Regel et al., 2010</u>); by manipulating prosodic style under controlled conditions, we extend this principle to the prosodic dimension of voice identity. Within this effect, for human voices, our observed late positivity aligns with speaker-specific communicative style violation effects; for AI voices, the observed late negativity that also signals expectation violations suggests that AI voice perception may resemble accented speech perception (<u>Jiang et al., 2020</u>). Third and most importantly, the observed comparable old/new effects across voice types indicate that identity perception mechanisms operate at a general level (<u>Giamundo et al., 2025</u>; <u>Zäske et al., 2017</u>); our findings extend this principle by demonstrating that such mechanisms transcend the biological vs. algorithmic origins of vocal signals.

Our findings also provide a theoretical basis for understanding AI voice agents in commercial applications: listeners do encode and remember AI speaker identities, suggesting that identity-related attributes may influence user acceptance and long-term engagement with synthetic voices (<u>Brandtzaeg et al., 2022</u>; <u>Jones et al., 2021</u>).

## Supporting information

bioRxiv_VoiceIDRecog_Supplementary_Materials

## Acknowledgements

We would like to thank Dr. Xiaolin Zhou for his encouragement, suggestions, and comments on this project’s experimental design, results interpretation, and presentation throughout WJC’s Master’s studies at Shanghai International Studies University, from the thesis proposal, through his academic writing course, to serving as thesis committee chair.

## Data and Code Availability

Behavioral data, EEG data, analysis code, and example stimuli are available at the Open Science Framework: https://osf.io/9b8pq/.

## Funding

This research was supported by the National Natural Science Foundation of China (Grant No. 32471109), awarded to X. Jiang. The PhD studentship of W. Chen was supported by a McGill-CSC (China Scholarship Council) Joint Scholarship, part of which is sourced from the Natural Sciences and Engineering Research Council of Canada (NSERC) Discovery Grant (RGPIN-2022-04363) awarded to M. D. Pell.

## CRediT

W. Chen: Conceptualization, Data curation, Formal analysis, Investigation, Methodology, Visualization, Writing – original draft. M. D. Pell: Resources, Supervision, Writing – review and editing. X. Jiang: Conceptualization, Funding acquisition, Investigation, Project administration, Supervision, Writing – review and editing.

## Declaration of generative AI and AI-assisted technologies in the manuscript preparation process

During the preparation of this work, the authors used Claude Sonnet 4.5 (Anthropic) to assist with highly customized data analysis and visualizations, as well as to refine wording in the manuscript. Grammarly was also used to check grammar and improve sentence clarity. After using this tool/service, the authors reviewed and edited the content as needed and take full responsibility for the content of the published article.

## Notes

https://osf.io/9b8pq/

### Competing Interest Statement

The authors have declared no competing interest.

