## Supplementary material for "Human and AI voice identities evoke shared neural signatures during speaker recognition across changes in speech content and prosody": bioRxiv_VoiceIDRecog_Supplementary_Materials

**Supplementary analysis 1: TF-MVPA reveals no significant decoding effects**

Time-frequency decomposition was performed using Morlet wavelet convolution in MATLAB (R2024b). Power was computed for 40 frequencies (4-80 Hz) using wavelets with cycles increasing linearly from 3 (at 4 Hz) to 10 (at 80 Hz). Baseline correction was applied using decibel conversion relative to pre-stimulus baseline (-500 to -250 ms), and data were downsampled by a factor of 3 along the time dimension. Multivariate decoding used linear discriminant analysis (LDA) via MVPA-Light toolbox (Treder, 2020). For each frequency-time point, classifiers distinguished between conditions using spatial power patterns across electrodes, evaluated with 5-fold cross-validation repeated 5 times (AUC metric). Group-level inference used cluster-based permutation testing (1,000 permutations, cluster-forming threshold $p < 0.05$, $\alpha = 0.05$).

For old vs. new speaker discrimination (based on correct judgments), decoding remained at chance across 4-80 Hz and 0-1000 ms for both human (mean AUC = 0.500 ± 0.054) and AI voices (0.501 ± 0.054). No significant clusters emerged for either voice type (0/6,680 pixels), with no difference between sources ($t(39) = -0.610$, $p = 0.545$). See **Figure S1**.

For prosodic consistency (same vs. different prosody among correctly recognized old speakers), performance similarly remained at chance for both human (0.500 ± 0.083) and AI voices (0.500 ± 0.084), with no significant clusters (0/6,680 pixels) and no voice source difference ($t(39) = 0.194$, $p = 0.847$). See **Figure S2**.

These null findings indicate that oscillatory power patterns do not reliably encode speaker identity or prosodic consistency information in the time-frequency domain.

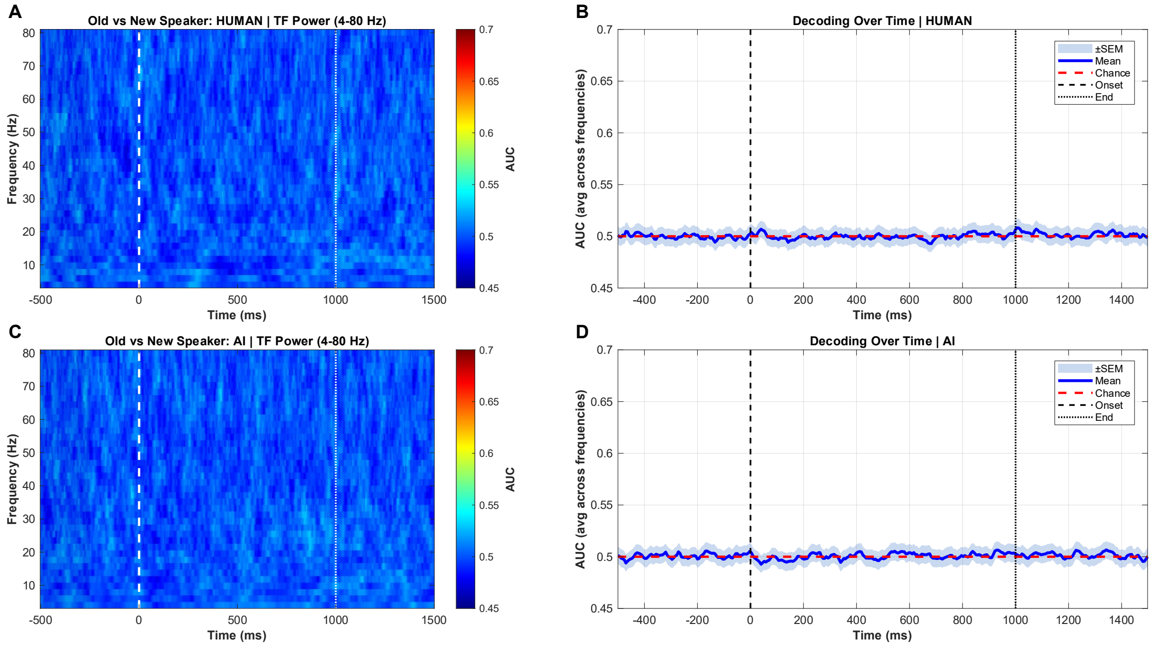

**Figure S1.** ***Time-frequency decoding of speaker identity (old vs. new).*** Multivariate decoding performance (AUC) for distinguishing correctly judged old from new speakers across frequencies (4-80 Hz) and time (0-1000 ms) for (A-B) human voices and (C-D) AI voices. Time-frequency maps (A, C) and frequency-averaged time courses (B, D) show no decoding above chance (AUC = 0.5, dashed line). No significant clusters identified (cluster-based permutation test, α = 0.05).

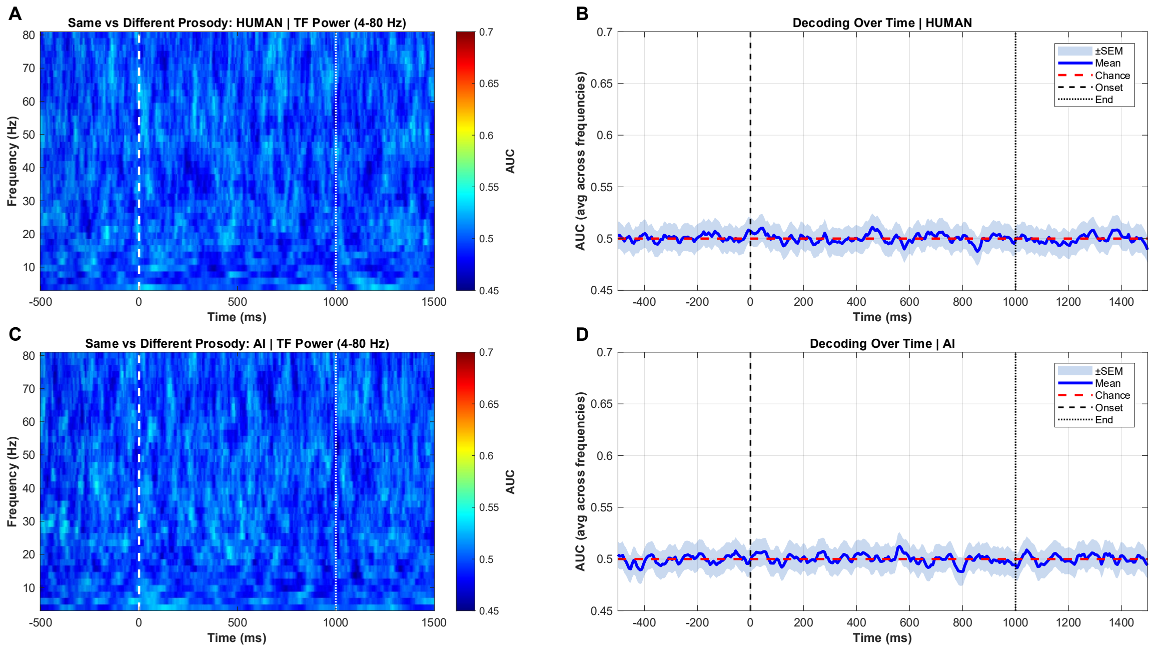

**Figure S2.** ***Time-frequency decoding of prosodic consistency (same vs. different).*** Multivariate decoding performance (AUC) for distinguishing same from different prosody among correctly recognized old speakers across frequencies (4-80 Hz) and time (0-1000 ms) for (A-B) human voices and (C-D) AI voices. Time-frequency maps (A, C) and frequency-averaged time courses (B, D) show no decoding above chance (AUC = 0.5, dashed line). No significant clusters identified (cluster-based permutation test, $\alpha = 0.05$).

### Supplementary tables

#### Table S1

*Linear mixed-effects model results for AI voices: Old vs. new speaker recognition (MVPA-identified windows)*

| Cluster | $F$ | $df$ | $p$ | | $\omega^2$ | $\beta$ | $SE$ | $\lvert t \rvert$ | $p$ | |
|---|---|---|---|---|---|---|---|---|---|---|
| cluster1 | 25.07 | 1, 7525 | <.001 | *** | 0.00319 | 1.340 | 0.267 | 5.01 | <.001 | *** |
| cluster2 | 16.38 | 1, 7540 | <.001 | *** | 0.00203 | 1.250 | 0.310 | 4.05 | <.001 | *** |
| cluster3 | 2.40 | 1, 7537 | 0.122 | ns | 0.00018 | 0.626 | 0.404 | 1.55 | 0.122 | ns |

*Note.* Three significant clusters identified via temporal decoding: 662-702 ms ($p = .029$), 758-844 ms ($p = .006$), and 866-1498 ms ($p < .001$). $\omega^2$ = omega-squared effect size; $\beta$ = regression coefficient; $|t|$ = absolute $t$-value. Positive $\beta$ indicates higher decoding accuracy for old speakers. **$p < .001$, *$p < .01$, *$p < .05$, ns = not significant.

#### Table S2

*Linear mixed-effects model results for human voices: Old vs. new speaker recognition (AI-identified windows)*

| Cluster | $F$ | $df$ | $p$ | | $\omega^2$ | $\beta$ | $SE$ | $\lvert t \rvert$ | $p$ | |
|---|---|---|---|---|---|---|---|---|---|---|
| cluster1 | 10.05 | 1, 7589 | 0.002 | ** | 0.00119 | 0.828 | 0.261 | 3.17 | 0.002 | ** |
| cluster2 | 2.80 | 1, 7592 | 0.094 | ns | 0.00024 | 0.489 | 0.292 | 1.67 | 0.094 | ns |
| cluster3 | 1.79 | 1, 7588 | 0.181 | ns | 0.00010 | 0.470 | 0.351 | 1.34 | 0.181 | ns |

*Note.* Human voices showed no significant clusters in temporal decoding analysis. Time windows borrowed from AI voice analysis for statistical testing: 662-702 ms ($p = .002$), 758-844 ms ($p = .094$), and 866-1498 ms ($p = .181$). $\omega^2$ = omega-squared effect size; $\beta$ = regression coefficient; $|t|$ = absolute $t$-value. Positive $\beta$ indicates higher decoding accuracy for old speakers. **$p < .001$, *$p < .01$, *$p < .05$, ns = not significant.

**Table S3**

*Linear mixed-effects model results for AI voices: Old vs. new speaker recognition*

| Window | Electrode | *F* | *df* | *p* | | ω² | β | *SE* | \|t\| | *p* | |
|---|---|---|---|---|---|---|---|---|---|---|---|
| N250 | Pz | 35.82 | 1, 309218 | <.001 | *** | 0.00011 | 0.29 | 0.05 | 5.99 | <.001 | *** |
| | P4 | 4.52 | 1, 308515 | 0.033 | * | 0.00001 | 0.08 | 0.04 | 2.13 | 0.033 | * |
| | C4 | 3.04 | 1, 308794 | 0.081 | ns | 0.00001 | 0.07 | 0.04 | 1.74 | 0.082 | ns |
| | P3 | 1.06 | 1, 308934 | 0.302 | ns | 0.00000 | -0.04 | 0.04 | 1.03 | 0.302 | ns |
| | C3 | 0.80 | 1, 308988 | 0.372 | ns | 0.00000 | 0.04 | 0.04 | 0.89 | 0.372 | ns |
| | Cz | 0.62 | 1, 308885 | 0.430 | ns | 0.00000 | 0.04 | 0.05 | 0.79 | 0.430 | ns |
| | F3 | 0.11 | 1, 308908 | 0.745 | ns | 0.00000 | -0.02 | 0.06 | 0.32 | 0.745 | ns |
| | F4 | 0.04 | 1, 309076 | 0.835 | ns | 0.00000 | -0.01 | 0.05 | 0.21 | 0.835 | ns |
| | Fz | 0.18 | 1, 309219 | 0.674 | ns | 0.00000 | -0.02 | 0.05 | 0.42 | 0.674 | ns |
| P300 | Pz | 137.83 | 1, 309411 | <.001 | *** | 0.00044 | 0.59 | 0.05 | 11.74 | <.001 | *** |
| | P3 | 47.15 | 1, 308966 | <.001 | *** | 0.00015 | 0.28 | 0.04 | 6.87 | <.001 | *** |
| | P4 | 45.46 | 1, 309072 | <.001 | *** | 0.00014 | 0.28 | 0.04 | 6.74 | <.001 | *** |
| | C3 | 12.06 | 1, 309130 | <.001 | *** | 0.00004 | 0.16 | 0.05 | 3.47 | <.001 | *** |
| | Cz | 6.37 | 1, 309257 | 0.012 | * | 0.00002 | 0.13 | 0.05 | 2.52 | 0.012 | * |
| | C4 | 2.68 | 1, 309363 | 0.102 | ns | 0.00001 | 0.07 | 0.04 | 1.64 | 0.102 | ns |
| | F3 | 0.23 | 1, 309060 | 0.631 | ns | 0.00000 | 0.03 | 0.06 | 0.48 | 0.631 | ns |
| | F4 | 0.76 | 1, 309283 | 0.382 | ns | 0.00000 | -0.05 | 0.05 | 0.87 | 0.382 | ns |
| | Fz | 0.17 | 1, 309449 | 0.682 | ns | 0.00000 | 0.02 | 0.06 | 0.41 | 0.682 | ns |
| LPC | Pz | 2715.60 | 1, 1518133 | <.001 | *** | 0.00178 | 1.35 | 0.03 | 52.11 | <.001 | *** |
| | P4 | 2364.50 | 1, 1517857 | <.001 | *** | 0.00155 | 1.02 | 0.02 | 48.63 | <.001 | *** |
| | P3 | 1761.10 | 1, 1517390 | <.001 | *** | 0.00116 | 0.88 | 0.02 | 41.97 | <.001 | *** |
| | Cz | 955.48 | 1, 1517675 | <.001 | *** | 0.00063 | 0.78 | 0.03 | 30.91 | <.001 | *** |
| | C4 | 867.51 | 1, 1517723 | <.001 | *** | 0.00057 | 0.66 | 0.02 | 29.45 | <.001 | *** |
| | C3 | 548.98 | 1, 1517294 | <.001 | *** | 0.00036 | 0.54 | 0.02 | 23.43 | <.001 | *** |
| | Fz | 247.42 | 1, 1517963 | <.001 | *** | 0.00016 | 0.44 | 0.03 | 15.73 | <.001 | *** |
| | F4 | 122.73 | 1, 1517178 | <.001 | *** | 0.00008 | 0.30 | 0.03 | 11.08 | <.001 | *** |

| | | | | | | | | | | | |
|---|---|---|---|---|---|---|---|---|---|---|---|
| | F3 | 38.85 | 1, 1516924 | <.001 | *** | 0.00002 | 0.19 | 0.03 | 6.23 | <.001 | *** |

*Note.* $\omega^2$ = omega-squared effect size; β = regression coefficient (estimated mean difference in amplitude, old – new) in µV; |*t*| = absolute *t*-value. First *p* and significance column refer to ANOVA main effect *F*-test; second *p* and significance refer to pairwise contrast. Positive β indicates more positive amplitudes for old speakers; negative β indicates more positive amplitudes for new speakers. **$p < .001$, *$p < .01$, *$p < .05$, ns = not significant.

**Table S4**

*Linear mixed-effects model results for human voices: Old vs. new speaker recognition*

| Window | Electrode | *F* | *df* | *p* | | ω² | β | *SE* | \|t\| | *p* | |
|---|---|---|---|---|---|---|---|---|---|---|---|
| | Pz | 48.93 | 1, 311628 | <.001 | *** | 0.00015 | -0.32 | 0.05 | 7.00 | <.001 | *** |
| | P3 | 40.70 | 1, 311890 | <.001 | *** | 0.00013 | -0.24 | 0.04 | 6.38 | <.001 | *** |
| | C3 | 28.83 | 1, 311860 | <.001 | *** | 0.00009 | -0.22 | 0.04 | 5.37 | <.001 | *** |
| | Fz | 12.38 | 1, 311921 | <.001 | *** | 0.00004 | -0.18 | 0.05 | 3.52 | <.001 | *** |
| N250 | C4 | 8.58 | 1, 311938 | 0.003 | ** | 0.00002 | -0.12 | 0.04 | 2.93 | 0.003 | ** |
| | P4 | 6.24 | 1, 311738 | 0.013 | * | 0.00002 | -0.09 | 0.04 | 2.50 | 0.012 | * |
| | Cz | 0.10 | 1, 311889 | 0.751 | ns | 0.00000 | -0.01 | 0.05 | 0.32 | 0.751 | ns |
| | F3 | 0.03 | 1, 311880 | 0.863 | ns | 0.00000 | 0.01 | 0.06 | 0.17 | 0.863 | ns |
| | F4 | 0.17 | 1, 311921 | 0.678 | ns | 0.00000 | -0.02 | 0.05 | 0.41 | 0.678 | ns |
| | Cz | 294.46 | 1, 311951 | <.001 | *** | 0.00094 | 0.83 | 0.05 | 17.16 | <.001 | *** |
| | C4 | 193.24 | 1, 311904 | <.001 | *** | 0.00062 | 0.60 | 0.04 | 13.90 | <.001 | *** |
| | C3 | 163.91 | 1, 311896 | <.001 | *** | 0.00052 | 0.57 | 0.04 | 12.80 | <.001 | *** |
| | F4 | 160.54 | 1, 311918 | <.001 | *** | 0.00051 | 0.66 | 0.05 | 12.67 | <.001 | *** |
| P300 | Fz | 136.27 | 1, 311930 | <.001 | *** | 0.00043 | 0.64 | 0.05 | 11.67 | <.001 | *** |
| | F3 | 134.07 | 1, 311837 | <.001 | *** | 0.00043 | 0.69 | 0.06 | 11.58 | <.001 | *** |
| | P4 | 32.22 | 1, 311780 | <.001 | *** | 0.00010 | 0.22 | 0.04 | 5.68 | <.001 | *** |
| | P3 | 22.68 | 1, 311890 | <.001 | *** | 0.00007 | 0.19 | 0.04 | 4.76 | <.001 | *** |
| | Pz | 4.36 | 1, 311777 | 0.037 | * | 0.00001 | 0.10 | 0.05 | 2.09 | 0.037 | * |
| | P4 | 1378.20 | 1, 1529408 | <.001 | *** | 0.00090 | 0.74 | 0.02 | 37.12 | <.001 | *** |
| | Pz | 370.31 | 1, 1529319 | <.001 | *** | 0.00024 | 0.48 | 0.02 | 19.24 | <.001 | *** |
| | P3 | 303.10 | 1, 1529579 | <.001 | *** | 0.00020 | 0.35 | 0.02 | 17.41 | <.001 | *** |
| LPC | Cz | 272.36 | 1, 1529560 | <.001 | *** | 0.00018 | 0.40 | 0.02 | 16.50 | <.001 | *** |
| | C4 | 183.63 | 1, 1529518 | <.001 | *** | 0.00012 | 0.29 | 0.02 | 13.55 | <.001 | *** |
| | F3 | 98.50 | 1, 1529316 | <.001 | *** | 0.00006 | -0.31 | 0.03 | 9.93 | <.001 | *** |
| | Fz | 44.93 | 1, 1529548 | <.001 | *** | 0.00003 | -0.18 | 0.03 | 6.70 | <.001 | *** |
| | F4 | 40.18 | 1, 1529480 | <.001 | *** | 0.00003 | -0.17 | 0.03 | 6.34 | <.001 | *** |

| | | | | | | | | | | |
|---|---|---|---|---|---|---|---|---|---|---|
| C3 | 14.13 | 1, 1529507 | <.001 | *** | 0.00001 | -0.09 | 0.02 | 3.76 | <.001 | *** |

*Note.* $\omega^2$ = omega-squared effect size; $\beta$ = regression coefficient (estimated mean difference in amplitude, old – new) in μV; $|t|$ = absolute $t$-value. First $p$ and significance column refer to ANOVA main effect $F$-test; second $p$ and significance refer to pairwise contrast. Positive $\beta$ indicates more positive amplitudes for old speakers; negative $\beta$ indicates more positive amplitudes for new speakers. **$p < .001$, *$p < .01$, $p < .05$, ns = not significant.

**Table S5**

*Linear mixed-effects model results for AI voices: Same vs. different prosody*

| Window | Electrode | *F* | *df* | *p* | | $\omega^2$ | β | *SE* | \|t\| | *p* | |
|---|---|---|---|---|---|---|---|---|---|---|---|
| Late (Negative) Prosody Effect | Pz | 910.16 | 1, 684902 | <.001 | *** | 0.00133 | 1.20 | 0.04 | 30.17 | <.001 | *** |
| | P3 | 415.27 | 1, 684486 | <.001 | *** | 0.00060 | 0.67 | 0.03 | 20.38 | <.001 | *** |
| | F4 | 196.42 | 1, 682677 | <.001 | *** | 0.00029 | 0.59 | 0.04 | 14.02 | <.001 | *** |
| | C4 | 152.71 | 1, 683909 | <.001 | *** | 0.00022 | 0.43 | 0.03 | 12.36 | <.001 | *** |
| | P4 | 130.58 | 1, 684568 | <.001 | *** | 0.00019 | 0.38 | 0.03 | 11.43 | <.001 | *** |
| | Cz | 114.86 | 1, 684303 | <.001 | *** | 0.00017 | 0.42 | 0.04 | 10.72 | <.001 | *** |
| | Fz | 98.06 | 1, 684344 | <.001 | *** | 0.00014 | 0.43 | 0.04 | 9.90 | <.001 | *** |
| | F3 | 95.36 | 1, 682273 | <.001 | *** | 0.00014 | 0.45 | 0.05 | 9.77 | <.001 | *** |
| | C3 | 23.33 | 1, 684377 | <.001 | *** | 0.00003 | 0.18 | 0.04 | 4.83 | <.001 | *** |

*Note.* Late Prosody Effect (500-900 ms) captures prosodic expectation violations. $\omega^2$ = omega-squared effect size; β = regression coefficient (estimated mean difference in amplitude, same – different) in μV; $|t|$ = absolute $t$-value. Positive β indicates more positive amplitudes for same prosody; negative β indicates more positive amplitudes for different prosody. **$p < .001$, *$p < .01$, *$p < .05$, ns = not significant.

**Table S6**

*Linear mixed-effects model results for human voices: Same vs. different prosody*

| Window | Electrode | *F* | *df* | *p* | | ω² | β | *SE* | \|t\| | *p* | |
|---|---|---|---|---|---|---|---|---|---|---|---|
| Late (Positive) Prosody Effect | C4 | 1252.50 | 1, 697828 | <.001 | *** | 0.00179 | -1.17 | 0.03 | 35.39 | <.001 | *** |
| | P4 | 934.66 | 1, 697779 | <.001 | *** | 0.00134 | -0.93 | 0.03 | 30.57 | <.001 | *** |
| | Cz | 816.96 | 1, 697826 | <.001 | *** | 0.00117 | -1.07 | 0.04 | 28.58 | <.001 | *** |
| | P3 | 527.43 | 1, 697844 | <.001 | *** | 0.00075 | -0.70 | 0.03 | 22.97 | <.001 | *** |
| | Fz | 434.09 | 1, 697747 | <.001 | *** | 0.00062 | -0.89 | 0.04 | 20.84 | <.001 | *** |
| | C3 | 428.29 | 1, 697823 | <.001 | *** | 0.00061 | -0.72 | 0.04 | 20.70 | <.001 | *** |
| | Pz | 385.43 | 1, 697838 | <.001 | *** | 0.00055 | -0.74 | 0.04 | 19.63 | <.001 | *** |
| | F4 | 375.42 | 1, 697703 | <.001 | *** | 0.00054 | -0.79 | 0.04 | 19.38 | <.001 | *** |
| | F3 | 18.45 | 1, 697694 | <.001 | *** | 0.00002 | 0.22 | 0.05 | 4.29 | <.001 | *** |

*Note.* Late Prosody Effect (500-900 ms) captures prosodic expectation violations. $\omega^2$ = omega-squared effect size; β = regression coefficient (estimated mean difference in amplitude, same – different) in µV; $|t|$ = absolute $t$-value. Positive β indicates more positive amplitudes for same prosody; negative β indicates more positive amplitudes for different prosody. **$p < .001$, *$p < .01$, *$p < .05$, ns = not significant.
